# Causal Mapping of Emotion Networks in the Human Brain: Framework and Preliminary Findings

**DOI:** 10.1101/214486

**Authors:** Julien Dubois, Hiroyuki Oya, J. Michael Tyszka, Matthew Howard, Frederick Eberhardt, Ralph Adolphs

## Abstract

Emotions involve many cortical and subcortical regions, prominently including the amygdala. It remains unknown how these multiple network components interact, and it remains unknown how they cause the behavioral, autonomic, and experiential effects of emotions. Here we describe a framework for combining a novel technique, concurrent electrical stimulation with fMRI (es-fMRI), together with a novel analysis, inferring causal structure from fMRI data (causal discovery). We outline a research program for investigating human emotion with these new tools, and provide initial findings from two large resting-state datasets as well as case studies in neurosurgical patients with electrical stimulation of the amygdala. The overarching goal is to use causal discovery methods on fMRI data to infer causal graphical models of how brain regions interact, and then to further constrain these models with direct stimulation of specific brain regions and concurrent fMRI. We conclude by discussing limitations and future extensions. The approach could yield anatomical hypotheses about brain connectivity, motivate rational strategies for treating mood disorders with deep brain stimulation, and could be extended to animal studies that use combined optogenetic fMRI.

## Introduction

How do networks of brain structures generate human emotions? Affective neuroscience has documented a wealth of data, primarily from activations observed in neuroimaging studies in response to emotional stimuli. This has provided us with an inventory of brain structures that participate in emotions, but little knowledge of their precise causal role. Studies in humans with direct electrical stimulation of structures such as the amygdala have shown causal links between brain regions and emotional responses, but these additional findings still leave us with scant knowledge of how emotions are implemented at the network level in the brain. The question is pressing for translational reasons as well. Deep-brain stimulation is being explored for a large number of neurological and psychiatric diseases, but with quite variable success. There are clear case studies of remarkable amelioration of depression, for instance—but only in some cases, limiting the generalizability of the results (Kennedy and al, 2011; Mayberg et al., 2005).

We think of emotions as functional, central brain states defined by their cause-and-effect relationships with other brain processes, and with stimuli and behaviors. Which stimuli reliably cause emotions? How do emotions in turn cause behavioral responses? And -- the topic of this paper --how do different brain regions causally interact with one another during emotion processing? The basic problem can be sketched in relation to the amygdala as schematized in **Figure 1**. The amygdala is activated by threat-related stimuli, lesions of the amygdala impair threat-related responses and (in humans) aspects of the experience of fear, and stimulation of the amygdala produces defensive behaviors (very roughly). Yet we do not understand the causal mechanisms that are responsible for these observations. Clearly, it is meaningless to say that "fear is in the amygdala”. The amygdala helps to orchestrate the many different causal effects of a fear state. But to understand these effects we need to map the causal relations between the amygdala and other brain regions, through which such effects are mediated. We know almost nothing about these causal relations in the human brain.

**Figure 1.**
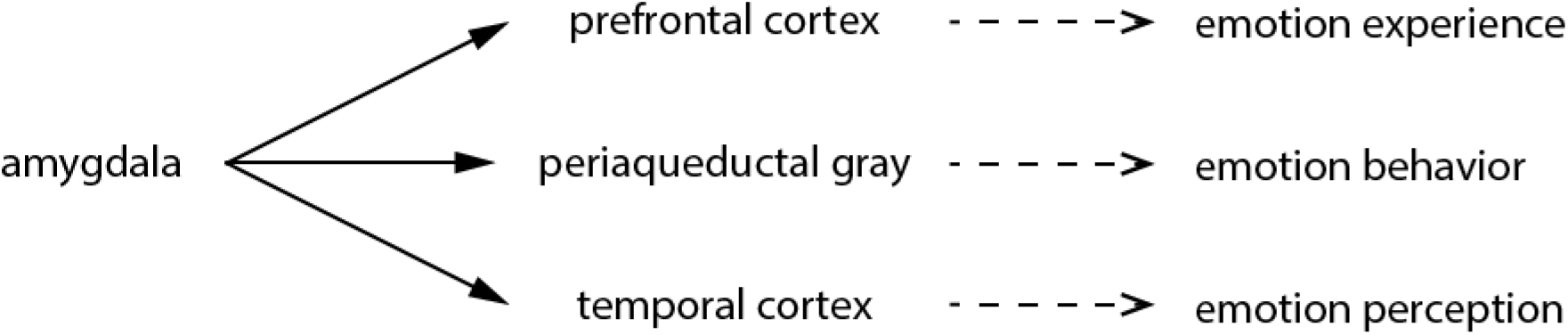
Amygdala connectivity with other structures. Schematized here as a very simplified causal graph are some of the main known interactions between the amygdala and cortical (prefrontal and temporal cortex) as well as subcortical (periaqueductal gray) structures, each of which in turn cause different components of an emotion that we can measure (which are hypotheses in this figure, just to make the conceptual point). Note that in the present paper we are omitting the periaqueductal gray for methodological reasons (insufficient spatial resolution in the parcellation scheme used for analysis of our MRI datasets), and we are focusing just on the brain networks, and not yet on the emotion measures (see **Figure 4**).

Studies in animals have begun to dissect the circuits responsible for processing emotion, and of course offer methodological tools that are unavailable in humans. For instance, experimental manipulation of brain activity in rodents and monkeys has provided insights into the causal roles of particular circuits, such as 3 the extended amygdala (Amaral and Adolphs, 2016; Shackman and Fox, 2016) and the hypothalamus (Lin et al., 2011). Electrophysiological measures permit single-neuron resolution in the recordings as well. And behavioral dependent measures, while they need to be interpreted carefully, have given us strong evidence for how these specific neuronal populations can cause specific emotional behaviors related to fear and aggression. One main limitation with these animal studies has been achieving a whole-brain field-of-view. Although specific circuits can be manipulated, e.g. through optogenetic or chemogenetic activation, the downstream effects are typically measured in only a very small subset of brain regions. One exciting future combination of methods is concurrent optogenetic stimulation with whole-brain fMRI (Lee et al., 2010; Liang et al., 2015), or with ultrasound imaging. This source of results will be a critical complement to the human studies in the future (see DISCUSSION). However, the homology to human emotions remains another main limitation (Adolphs and Anderson, 2018).

Elucidating the causal networks that underlie emotion processing is one of the most important but also most difficult challenges faced by affective neuroscience. It is important because only an account at the level of causal mechanisms can really explain brain processing, and because only such an account can yield insights that allow us to manipulate brain function (for instance, with interventions aimed to treat mood disorders). Yet it is difficult because almost all data from the human brain are correlational in nature, making it unclear how to infer causality from typical neuroimaging and electrophysiological studies. Here we demonstrate the promise of a new technique - concurrent electrical stimulation and fMRI - and a new method in causal discovery - the fast greedy equivalence search - to obtain large-scale causal models that describe how different brain regions interact. We begin by briefly reviewing some of the findings from affective neuroscience, with an emphasis on the amygdala, and then outline the logic of causal discovery, before presenting our approach and pilot data to support it.

### Emotion and the amygdala

Data from lesion studies and fMRI in humans, and from a range of approaches in animals, consistently implicate the amygdala (**Figure 2**), the medial prefrontal cortex, the insula, the hypothalamus, and the periaqueductal gray in emotions. These structures function as components of considerably more distributed systems, and attempts to localize particular emotions (fear, sadness, etc.) to any one of these structures have been largely unsuccessful (Lindquist et al., 2012), even though specific emotions can be classified from distributed activation patterns in neuroimaging studies (Kragel and LaBar, 2015; Nummenmaa and Saarimäki, 2017; Saarimäki et al., 2015; Wager et al., 2015). While debates about how to interpret the data thus far remain (Adolphs, 2017a, b; Barrett, 2017a, b), neuroimaging and electrophysiological findings have supported a picture of emotion states implemented by distributed cortical and subcortical circuits. A large open question is thus how to understand the functional role played by specific brain structures: what exactly does a structure such as the amygdala contribute to processing fear (or any other emotion) and at what point in the distributed processing of that emotion does it exert meaningful causal effects on the other components of the network?

**Figure 2.**
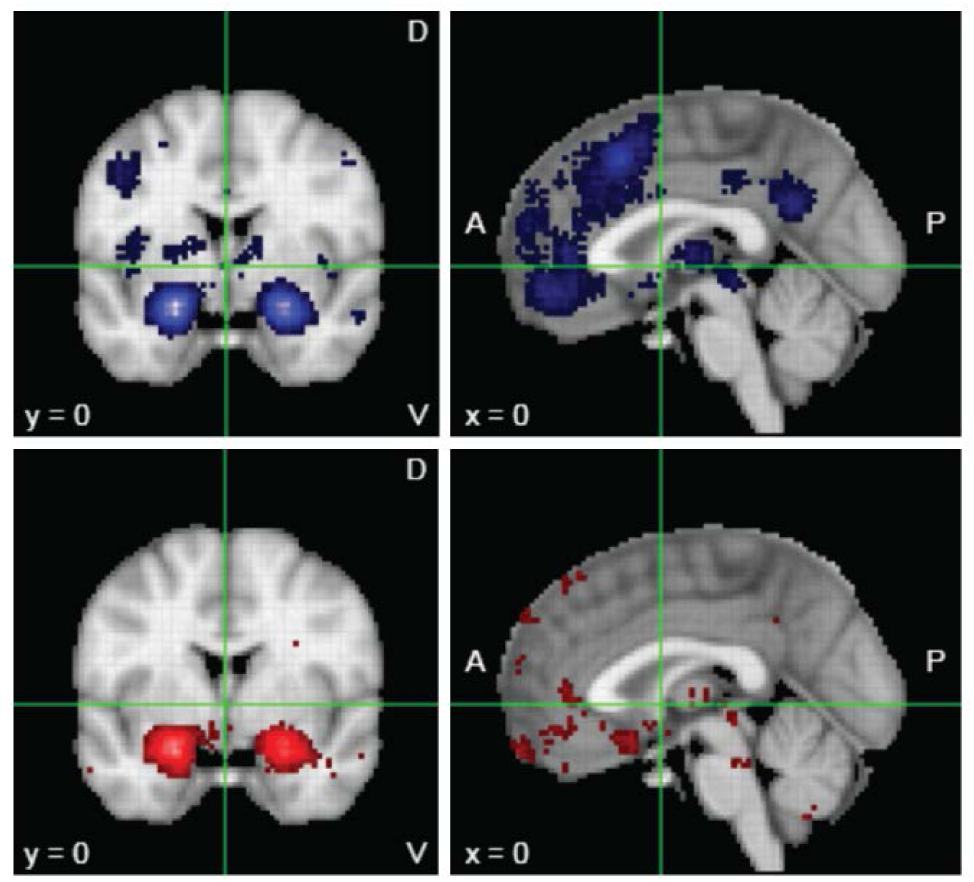
Meta-analytic mapping of brain activations for emotion. Neurosynth maps for the keyword “emotion” yielded results from 790 fMRI studies (*www.neurosynth.org*). In the top panels (in blue) are the “forward inference” maps (all activations above a statistical threshold having to do with “emotion”). However, most or all of these regions are also activated in many other studies that do not have anything to do with emotion. A more specific analysis would ask which regions were activated only by those studies containing the keyword “emotion", and not in studies that did not contain the word "emotion". This “reverse-inference" map is shown in the bottom panels (in red). Both analyses highlight the prevalence of reporting the amygdala, and to some extent the prefrontal cortex. This large bias in the literature has resulted in a strong belief that the amygdala is causally involved in emotion, a conclusion that is not yet warranted from the extant data.

The amygdala together with the bed nucleus of the stria terminalis (Fox and Shackman, 2017) appears to serve a role as an organizing center in these circuits, coordinating the multiple cognitive, autonomic, and behavioral effects of an emotion (Davis, 1992; Whalen and Phelps, 2009). Both the known structural connectivity of the amygdala (in monkeys and rodents) and structure-function relationships in particular tasks, such as Pavlovian fear conditioning, strongly support this view (**Figure 3**). Yet the direct evidence for causal relationships is nearly nonexistent in humans, and remains very sparse even in animals.

**Figure 3.**
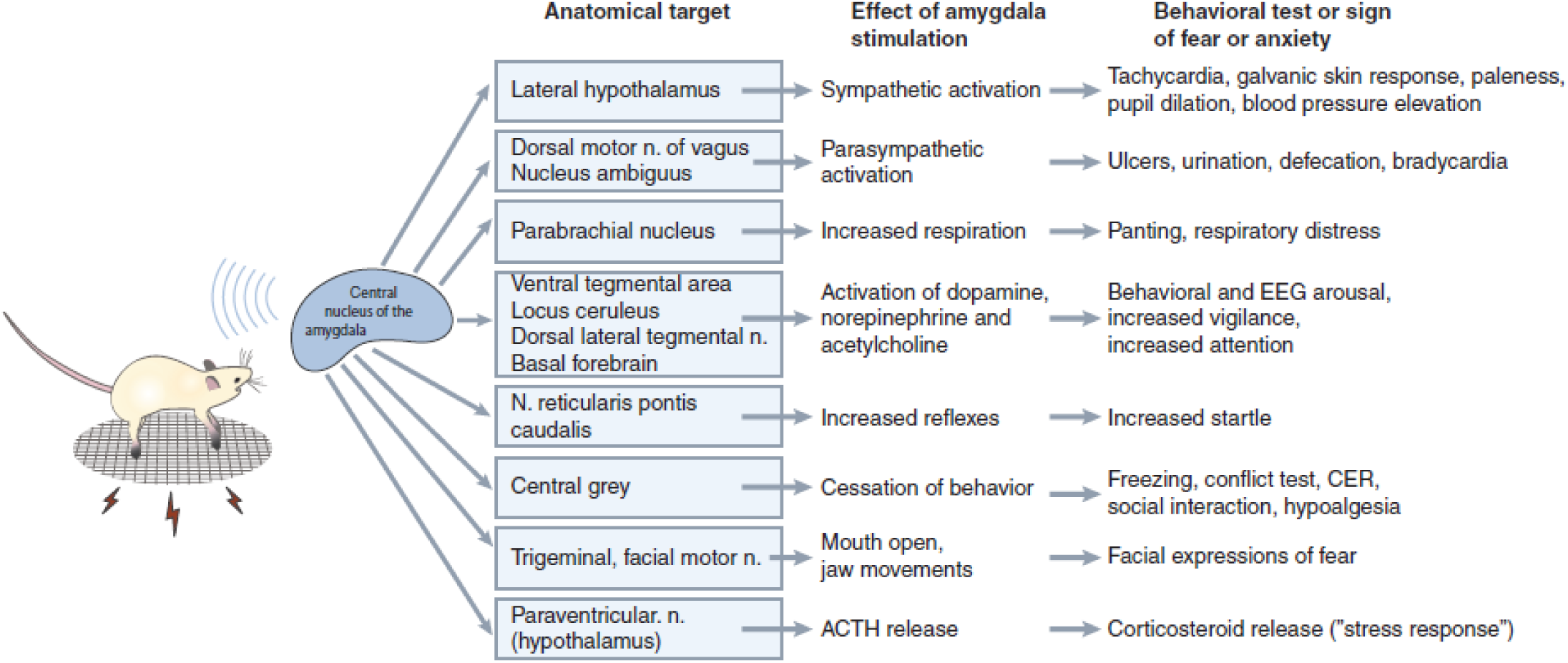
Connectivity of the central nucleus of the amygdala. The illustration summarizes the function of this amygdala nucleus in coordinating emotion components through its multiple causal effects on other brain regions. Note that the picture is in fact considerably more complicated, since the central amygdala also closely interacts with the adjacent bed nucleus of the stria terminalis in triggering the target effects shown here (Shackman and Fox, 2016). Reproduced from (Davis, 1992).

While lesion studies argue for the necessary role of a brain structure, they do not elucidate the neural mechanisms through which the lesioned tissue contributes to normal function. Lesions of the amygdala, in animals as well as humans (Amaral and Adolphs, 2016) result in impairments in fear processing. In humans, these can include strikingly selective deficits in the recognition of fear from facial expressions (Adolphs et al., 1994) and in the conscious experience of fear to exteroceptive threats (Feinstein et al., 2011) but not to certain interoceptive stimuli (Feinstein et al., 2013). However, studies in monkeys have shown that the causal consequence of an amygdala lesion on the brain is extremely complex, including widespread network changes in many other regions (Grayson et al., 2016). So although amygdala lesions have some effect on emotional behaviors and conscious experience, understanding the causal mechanisms explaining this effect requires additional measures. Indeed, some current theories of the conscious experience of emotion argue that the amygdala’s role in feelings and emotions is mediated entirely through cortical structures (LeDoux and Brown, 2017). Similarly, lesions of the ventromedial prefrontal cortex (vmPFC) can lead to alterations in emotional behavior, such as impaired autonomic responses (Bechara et al., 1996), dysregulation of anger (Koenigs and Tranel, 2007), and atypical moral judgment (Koenigs et al., 2007). Once again, it is difficult from this to infer the causal mechanisms, which may involve additional brain regions with which the vmPFC is connected. For instance, lesions of the vmPFC result in abnormal activation of the amygdala when lesion patients undergo fMRI (Motzkin et al., 2015).

Thus almost all of the evidence for the causal mechanisms behind emotions is very indirect and tenuous. It derives from a combination of structural connectivity studies in animals (the basis for most of **Figure 3**), piecemeal assembly of evidence across very different studies in the literature (much of the basis of **Figure 2**), or flawed inference of causation from correlation (nearly everything based on neuroimaging alone, or electrophysiology alone). While this problem is well known, it is also well ignored, in the hope that sheer accumulation of correlative data of various kinds could in and of itself provide us with an understanding of the causal mechanisms.

Here we describe a research program that could take us from these heterogeneous beginnings to a principled approach for investigating causal architecture for emotions. Which brain regions are involved, how are they causally related to one another, and how do they in turn cause particular components of emotions? We suggest (and will show below) that parts of this broad and ambitious aim can in fact already be addressed with human neuroimaging data alone, using a novel causal discovery approach which we detail here. Parts of the aim also require new methods that make possible direct causal perturbation in the human brain, which we have recently developed (Oya et al., 2017). We provide initial results and a workflow for generating causal models from both of these types of data.

### Causal Discovery

The range of methods for analyzing brain function at the network level vary from essentially descriptive (such as looking at correlations between regions, i.e., standard functional connectivity approaches) to methods for inferring parameters of pre-specified models (such as dynamic causal modeling (DCM) (Friston, 2011; Friston et al., 2013)). All of these have tradeoffs: standard “functional connectivity” from correlations does not provide a causal model; DCM is limited by our knowledge of the physiological basis of the BOLD response, the lack of a search algorithm over models, and poor scalability. In its classical implementation, for instance, DCM can only test very simple models (10 - 20 nodes) that are too restrictive for understanding realistic whole-brain networks (Smith, 2012; Smith et al., 2011). It should be noted that there are continuous improvements, such as the novel regression-DCM approach, which has been scaled to 66 regions with 300 free parameters (Frässle and al, 2015; Frässle and al., 2017). But the primary challenge for DCM remains unsolved: how does one constrain reverse inference and search efficiently over candidate models given measured data? The number of candidate DCM models is enormous even for a small number of nodes, and there exists no efficient algorithm to search over all of them. The sweet spot, we believe, lies in the middle between standard functional connectivity approaches and physiologically-based models like DCM: causal modeling that takes advantage of the strengths of current BOLD-fMRI, and that enables the integration of experimental and observational data.

Causal models can be thought of as generative models that make predictions about what we might observe and about how we might achieve certain effects through experimental manipulation (e.g., brain stimulation to treat a mood disorder) (Pearl, 2009; Spirtes et al., 2000b). The usual way to depict the causal relationships between variables (causes from one brain region to others, in our case) is with a drawing called a causal graph. **Figure 3** above could be interpreted this way: the amygdala causes effects in the brainstem and hypothalamus which in turn cause effects in brain and behavior that are our dependent measures. Direct causal connections are taken to be relative to the set of variables depicted, thus a chain of three variables A→B→C without a direct arrow from A to C would indicate that B causally screens off A from C, that is, B completely mediates the causal effect of A on C. In terms of brain structures, we would think of this as region A providing inputs to B which in turn provides inputs to C, but without any direct connections from A to C.

Temporal order is often taken to be fundamental to causality, but for causal discovery it can be misleading. Intuitively, we would expect the cause A to precede its effect B, but if both A and B are effects of a common cause C, then B may well take longer to manifest itself than A, giving the impression as if A caused B. For causal discovery from fMRI data, actual temporal order of action potential events cannot be resolved, since these operate at a millisecond range that exceeds the temporal resolution of hemodynamic measures. Moreover, if one considered temporal order (which some analyses of BOLD-fMRI indeed attempt; (Friston et al., 2013)), one must take into account the interactions between the sampling rate and the rate at which the true underlying process operates, as well as regional differences in hemodynamic coupling. Appropriate temporal resolution and the absence of unmeasured common causes are the key assumptions underlying a valid inference to causal relations using Granger Causality (Granger, 1969). Concerns about these, among others, suggest that - though it has been used with fMRI data - Granger Causality is not well-suited for the causal analysis of fMRI (Smith 2011)(Stokes and Purdon, 2017).

Instead, modern causal discovery algorithms disregard temporal order and use the independence structure observed in the data in order to infer the underlying causal structure (see (Eberhardt, 2017) for an accessible brief review). The general idea of using the independence structure for causal inference goes back to the Principle of Common Cause (Reichenbach, 1956): if two variables are dependent, then either one causes the other, or vice versa, or there is a common cause of the two variables. Conversely, if two variables are independent, then they cannot (in general) be causes of one another or effects of a common cause. The independence and dependence structure found in data can thus be used to constrain candidate causal models that would explain the observed data.

The simplest case in which a fully oriented causal structure can be uniquely determined from observational data is when there are three variables A, B and C, and A and C are probabilistically independent, but A and B, and C and B are dependent. If there are no unmeasured confounders, then this independence structure provides a signature that uniquely identifies A→B←C as the causal graph connecting the variables. More generally, it is well understood how to use the observed independence structure over the variables to constrain the underlying causal structure. Even if the causal structure cannot be uniquely identified, the set of equivalent causal structures can be identified (the equivalence class consists of all causal structures consistent with the observed data). Our results below demonstrate the power of this approach, using a new variant of a causal discovery algorithm (Ramsey et al., 2016) that scales to large sets of resting-state fMRI data (here, the Human Connectome Project dataset and the MyConnectome Project dataset). Our variables (the nodes of the causal graph; A,B,C etc.) will be the brain regions into which a whole brain is parcellated, and whose causal relations are the question of interest.

It is worth contrasting the results of the causal inference algorithm with purely correlation-based techniques commonly applied to resting-state fMRI data. If the true causal structure has the form A→B←C, then the Pearson correlation matrix will have all non-zero entries except for the correlation of A and C. However, the inverse correlation matrix will have no non-zero entries, since all partial correlations conditional on all remaining variables are non-zero; in particular A is not independent of C given B. However, if the true causal structure is A→B→C, then all Pearson-correlations will be non-zero, but the partial correlation of A and C given B is zero. As the two simple examples above indicate, an adjacency structure based on either the Pearson correlation matrix, or on the inverse correlation matrix (partial correlation), is not a representation of the causal structure. Causal inference algorithms disentangle exactly what inferences can be drawn about the presence and absence of causal connections from the independence structure (see also **Figure 7**).

Another way of inferring causal models is through experimental intervention. Whereas the approach above relies on conditional probabilities that are merely observed, such as analysis of resting-state fMRI data, experimental intervention corresponds to the conditional probabilities produced through what Judea Pearl coined as the “do” operator (Pearl, 2009). This is the type of data often sought in animal studies of emotion, for instance through optogenetic manipulation of the activity of a node (brain region) with full or partial experimental control. This concept also underlies the essence of randomized controlled trials: by randomizing subject assignment to treatment group, one is experimentally intervening and (in the large sample limit) breaking all confounding causes (Fisher, 1990). Once again, we can think of wanting to infer A→B in a brain (with A and B as distinct regions of interest) where there are many other possible causes at work. This time we experimentally activate A to see if we observe a change in B. To the extent that the activation fully controls A, this experimental manipulation amounts to breaking all the causal effects that could act on A (all the arrows going into A), in particular those that might act as confounders in an observational setting. As we will show in our R**ESULTS** below, in humans this can now be accomplished through the rare opportunity of electrically stimulating the amygdala while measuring whole-brain fMRI. While the electrical stimulation will not fully control the pattern of activation, it can still help us to infer causal relationships with respect to the amygdala: It can help confirm that the relations detected in the observational dataset (such as resting-state fMRI) are not spurious, and possibly even help to orient causal arrows that could not be oriented from merely observational data.

In sum, the particular causal discovery algorithm that we used takes the observational (e.g., resting-state fMRI) data and tries to identify a sparse graph structure that accounts for all of the dependencies (the standard Pearson correlation matrix of the data). Our subsequent direct electrical stimulation experiments provide a completely different, interventional, set of data for comparison, validation, and possible further inference and refinement of the causal graph. The final result leverages at least three applications. First, it can be used to generate hypotheses, and interpret data, from many fMRI studies: studies that use emotional stimuli to produce brain activations, for example, could now be interpreted in terms of a causal mechanism implemented among the brain regions that instantiates an emotion state. Second, the results could be used to make neuroanatomical predictions: in their strongest interpretation they imply actual anatomical connectivity corresponding to the edges in the graph. And third, they make predictions about the effects of interventions through deep brain stimulation—predictions that could be used for strategically guided treatment of mood disorders. All of these are future goals of the framework we present here. The METHODS below outline our overall approach, the RESULTS provide preliminary findings, and the DISCUSSION notes the limitations and hurdles still remaining to be solved (cf. **Figure 4**).

**Figure 4.**
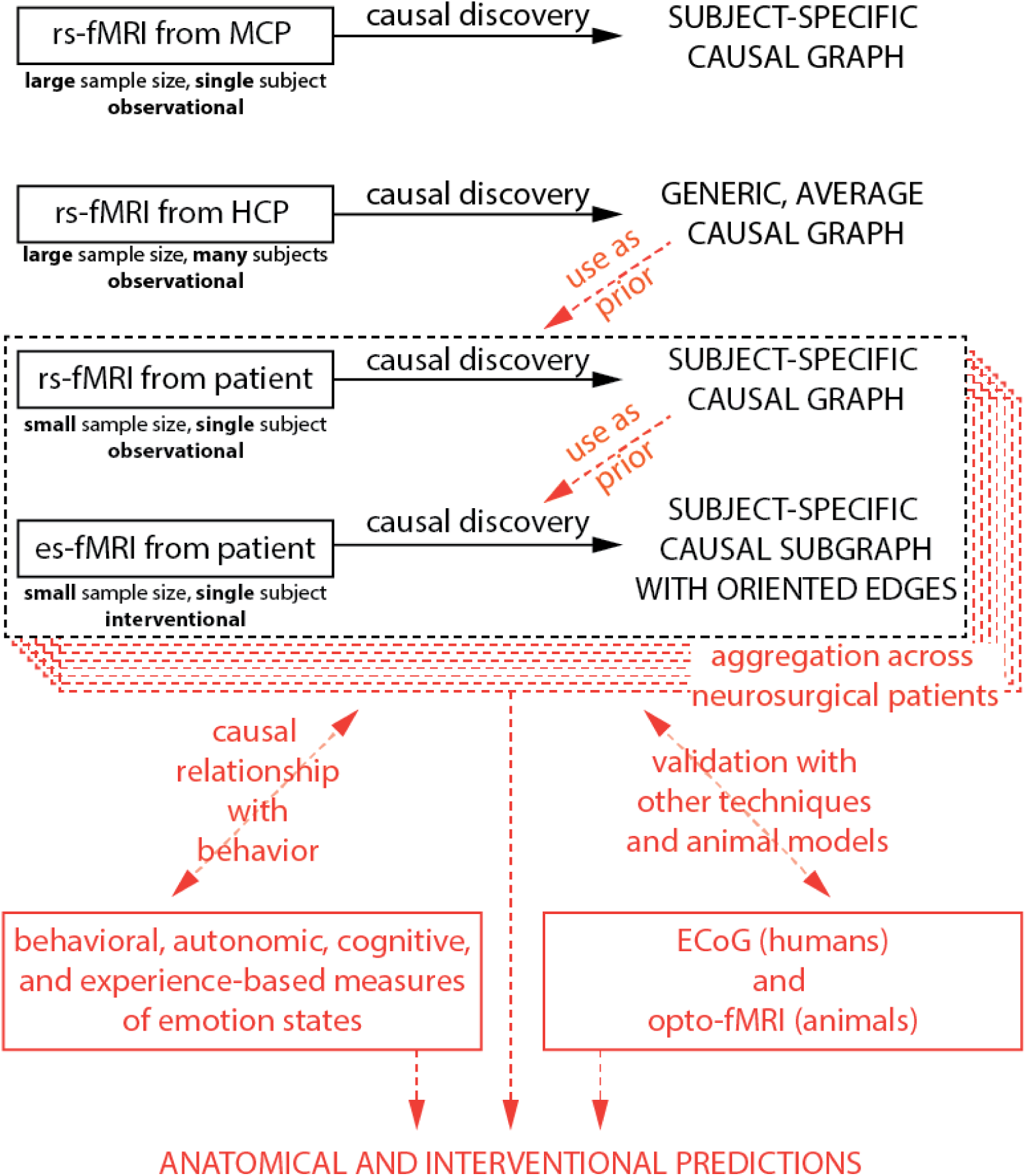
Proposed framework for causal analysis of emotion networks. In black, analyses presented in this manuscript; in red, future extensions outside the scope of the present paper. Abbreviations: HCP: Human Connectome Project dataset. MCP: MyConnectome Project dataset. rs-fMRI: resting-state fMRI. es-fMRI: concurrent electrical stimulation with fMRI.

**Figure 5.**
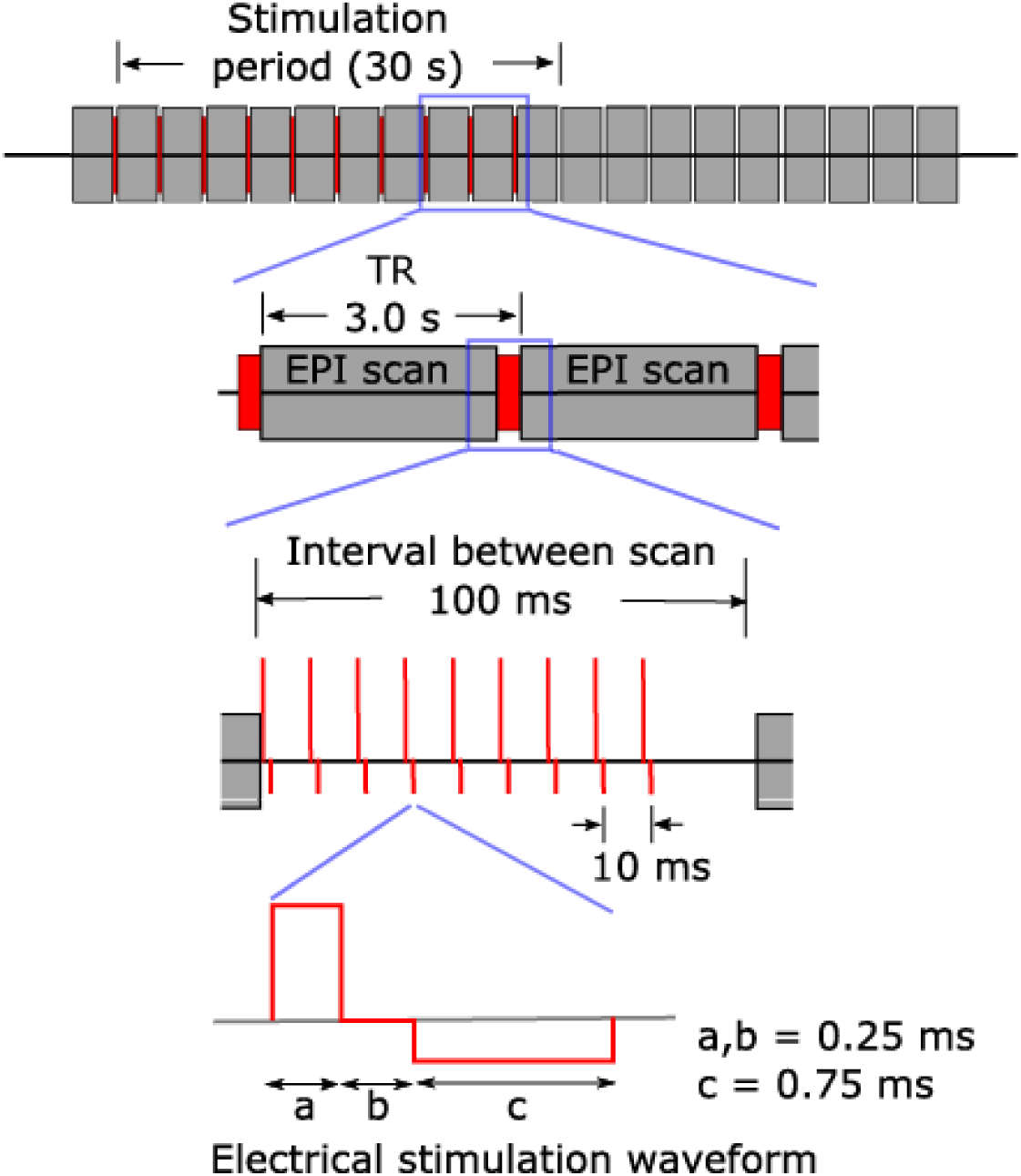
Electrical stimulation with concurrent fMRI. Reproduced from (Oya et al., 2017). es-fMRI protocol used. Each gray block is one whole-brain fMRI volume, red is the electrical stimulation shown at increasing magnification from top to bottom. Electrical stimuli were delivered to the subjects between EPI volume acquisition, during a 100 ms blank period, ensuring no temporal overlap with RF transmission nor with gradient switching. Modified charge-balanced constant-current bi-phasic pulses were used.

## Methods

Our approach is roughly hierarchical in nature (**Figure 4**) and consists of three main steps. These three steps are based on the analysis of existing resting-state data from large databases, comparison to resting-state data from individual neurosurgical patients, and comparison with the patient’s graph using direct electrical stimulation-fMRI data from that patient. Future extensions beyond the scope of the present paper include (in red): formally integrating information from the large datasets as prior constraints for analysis of each patient’s data; comparing, and combining, results across multiple patients; and leveraging the results into hypotheses that could be tested with additional experiments and additional methods—hypotheses about anatomical connectivity, about individual differences, and about clinical intervention efficacy.

### Step 1: Causal inference from observational resting-state fMRI data with a large number of samples

We first searched for causal graphs using observational data alone, taking advantage of large publicly available, high quality datasets.

#### Datasets

We used data from two public repositories, the 1200 subjects release of the Human Connectome Project (HCP) (Van Essen et al., 2013) and the MyConnectome Project (MCP) (Laumann, 2015; Poldrack, 2015). Both of these feature resting-state fMRI data with a large number of samples. They yield largely complementary information: the HCP provides data from almost 1200 subjects, which we can combine to de-emphasize individual-subject idiosyncrasies and thus extract a network structure with the highest level of generalizability; by contrast, the MCP data comes from a single subject scanned almost 100 times over the course of one year and allows us to derive a very reliable causal graph at the single subject level, with inevitable idiosyncrasies compared to the group-level graph derived from the HCP data.

Acquisition parameters and preprocessing of the resting-state fMRI data in both projects are described in their respective original publications (HCP, (Glasser et al., 2013); MCP, (Poldrack, 2015)). Briefly, in the HCP dataset, each subject underwent two sessions of resting-state fMRI on separate days, each session with two separate 15 minute acquisitions generating 1200 volumes (customized Siemens Skyra 3 Tesla MRI scanner, TR = 720 ms, TE = 33 ms, flip angle= 52°, voxel size = 2 mm isotropic, 72 slices, matrix = 104 × 90, FOV = 208 mm x 180 mm, multiband acceleration factor = 8). The two runs acquired on the same day differed in the phase encoding direction (left-right and right-left), which leads to differential signal intensity especially in ventral temporal and frontal structures. We used only the right-left phase encoding dataset, since this optimized signal in the right medial hemisphere, enabling the best comparisons to the electrical stimulation of the right amygdala in the neurosurgical patient #384 we describe further below. For the MCP dataset, a single subject underwent one 10 minute resting-state run, generating 518 volumes in each of 89 “production” sessions acquired on separate days (Siemens Skyra 3-Tesla MRI scanner, TR = 1,160 ms, TE = 30 ms, flip angle = 63°, voxel size = 2.4 mm x 2.4 mm x 2.0 mm, 68 slices, inter-slice distance factor = 20%, axial-coronal tilt 30 degrees above AC/PC line, matrix = 96 × 96, FOV = 230 mm FOV, multiband acceleration factor = 4). The HCP data was downloaded in its minimally preprocessed form, i.e. after motion correction, B0 distortion correction, coregistration to T_1_-weighted images and normalization to MNI space; while the MCP data was downloaded in its fully processed form (in native space), including minimal preprocessing and followed by resting-state specific denoising as described in (Laumann, 2015): censoring of frames with framewise displacement > 0.25 mm; regression of signals from whole brain, white matter and ventricles and their derivatives; regression of 24 movement parameters derived by Volterra expansion; and bandpass filtering 0.009 < f < 0.08 Hz. For consistency with MCP data, we replicated the same denoising pipeline (minus frame censoring) in Python (v2.7) and applied it to the minimally preprocessed data from the HCP.

#### Brain parcellation

The first step in a network analysis of the brain is the definition of its nodes (Sporns, 2013). There are two main approaches to defining network nodes in the brain: nodes may be a set of overlapping, weighted masks, e.g. obtained using independent component analysis (ICA) of BOLD fMRI data (Smith et al., 2013); or a set of discrete, non-overlapping binary masks, also known as a hard parcellation. Hard parcellations come in many flavors, in terms of what data they are based on (anatomical data only, functional data only, or multi-modal), and whether they are group-based or individually-derived. Parcellating the brain is an area of intense investigation, and significant progress has been made in recent years (Glasser et al., 2016; Gordon et al., 2014). We eventually aim to utilize the most recent developments in surface-based analysis and multi-modal surface matching (MSM) (Robinson et al., 2014), which divides the brain into about 400 regions. (Indeed, a future aim would be to use voxelwise data for the analyses we describe below, and to use causal discovery results to aggregate these into larger parcels based on their causal connectivity). However, applying these methods to patient data with implanted electrodes is difficult because MSM requires several types of data (such as high-resolution anatomical scans for precise surface reconstruction, field maps for precise co-registration of functional and anatomical data, and companion task and rest scans), which are not always available due to clinical constraints. Here, in this proof-of-concept paper, we used a more common volumetric parcellation in MNI space that divides the brain into 110 regions, based on the classical Harvard-Oxford anatomical atlas <u>(http://www.cma.mgh.harvard.edu/fsl_atlas.html)</u>. Specifically, we derived maximum probability labels from the probabilistic Harvard-Oxford cortical and subcortical atlases distributed with FSL, using a 25% probability threshold for label assignment. We omitted one parcel from the Harvard-Oxford atlas, the brainstem, since this treats the entire region as a single functional object, a parcellation that we felt was too coarse for our purposes. See **Supplementary Table 1** for the complete list of parcels we used.

#### Timeseries Extraction and Data Selection

HCP data was already transformed to MNI space (the same space as the Harvard-Oxford parcellation that we used) by the minimal preprocessing pipeline (Glasser et al., 2013). We derived a gray matter mask for each subject using information in ribbon.nii.gz and wmparc.nii.gz (from the individual HCP MNINonLinear directories), and restricted Harvard-Oxford parcels to gray matter voxels for each subject. For MCP data, MNI-space Harvard-Oxford labels were warped to the individual space via the MNI152 T1w template and the coregistered average of all 15 MCP T1w anatomical scans (Freesurfer 5.3.0 recon-all pipeline), using a diffeomorphic symmetric normalization (SyN) warp as implemented in ANTs 1.2.0 (antsRegistrationSyN.sh script). Similarly, the warp from the MCP T1w average to a mean MCP EPI template was estimated and applied to the Harvard-Oxford labels using ANTs (the N-back EPIs were used to construct the mean EPI template, in place of the rsBOLD EPIs, because the latter were global mean-subtracted and therefore unusable for registration; this is possible because all MCP EPIs are transformed to the same space in the downloaded data). The gray matter mask output from the Freesurfer pipeline was used in combination with the individual-space Harvard-Oxford atlas labels to restrict ROIs to gray matter voxels. Timeseries for all Harvard-Oxford parcels were extracted using the 3dROIstats function provided by AFNI 17.2.04 (Cox, 2012).

We generated three datasets from the HCP and MCP data:

*Dataset #1:* HCP sparsely sampled (HCPs). For this dataset we chose samples sparsely from two resting-state fMRI runs with right-left phase encoding (rfMRI_REST 1_RL and rfMRI_REST2_RL) from each of 880 unique subjects (with complete datasets and relative movement root-mean-square less than 0.15mm for both runs; see subject list in **Supplementary Table 2**a), taking one volume every 35^th^ TR (i.e. every 25.2s). This resulted in 68 samples per subject, for a total of 59,840 samples. We split this dataset into 11 completely non-overlapping subsets (80 subjects each) for “horizontal” reliability analysis, yielding 5,440 samples per subset. This provides the most general dataset with a large number of samples, pooling over a large number of subjects for representativeness, and sampling sparsely to eliminate autocorrelation and thus maximize statistical independence between samples. Representativeness across subjects here trades off with the possibility of introducing additional dependencies in the data due to the pooling of data from multiple subjects. The next dataset therefore focuses on data from a single individual.

*Dataset #2:* MCP sparsely sampled (MCP_s_). This dataset chose samples sparsely from 80 sessions of the MCP (see session list in **Supplementary Table 2**b), taking one volume every 22^nd^ TR (i.e. every 25.5s). This resulted in 23 samples per session, for a total of 1,840 samples. We generated 20 non-overlapping datasets for reliability analysis, by shifting the starting volume (note that these datasets are not completely independent from one another due to autocorrelation of the fMRI BOLD signal).

*Dataset #3:* MCP densely sampled (MCP_d_). This dataset chose all volumes from 80 sessions in the MCP yielding a total of 41,440 samples. We split this dataset into 8 non-overlapping subsets (10 sessions each) for horizontal reliability analysis, yielding 5,180 samples per subset.

**Causal discovery algorithm**

We used a version of the Fast Greedy Equivalence Search (FGES) algorithm for causal discovery (Ramsey et al., 2016), a variant of the better known Greedy Equivalence Search (Chickering, 2002) that was optimized to large numbers of variables. The algorithm takes as input measurements over a set of variables that one can think of as nodes in the causal graph, in this case the mean BOLD signal obtained for each region of interest in a parcellated human brain (the 110 parcels provided by the Harvard-Oxford atlas). FGES produces as output causal graphs that describe inferred direct causal connections between any pair of brain regions (the adjacency matrix), and, where possible, the orientations of these causal effects, i.e. whether brain region A causes BOLD response in brain region B, or vice versa. For any specific output causal graph, one can also estimate the strength of each causal connection (the effect size of each edge) using a linear Gaussian model. We next describe the algorithm in more detail.

The algorithm is a greedy optimization algorithm that operates in two phases, a forward phase in which edges are added to the graph, and a backward phase in which edges are removed. Under the assumption that the true causal model is causally sufficient (there are no unmeasured common causes), acyclic (there are no feedback cycles) and that the data is independent and identically distributed and not subject to sample selection bias, the FGES algorithm returns in the infinite sample limit the Markov equivalence class of the true causal structure with probability 1. That is, as sample size increases, the output of FGES converges towards (a representation of) a set of causal structures that are not distinguishable from the true causal structure on the basis of their probabilistic independences. For example, the three causal graphs A→B→C, A←B→C and A←B←C together form a Markov equivalence class under the given assumptions, since in all three structures A is independent of C given B, but no other (conditional) independences hold among the variables. Without experimental interventions or further assumptions (e.g. concerning time order or the parametric form) these three causal structures cannot be distinguished. In contrast, (as noted earlier in this paper) the causal structure A→B←C has a unique independence structure --it only satisfies that A and C are marginally independent, but no conditional independence -- and therefore forms a singleton Markov equivalence class.

The central idea underlying the FGES algorithm is the insight that one can construct a score tracking the posterior probability of a causal graph given a dataset such that the score is (i) decomposable into local scores for each edge, and (ii) gives the same value for graphs that are Markov equivalent. This insight provides the basis for a greedy search method that starts with an empty graph over the set of variables and then at each stage determines *locally* whether an edge addition improves the *global* score over the current causal graph. The edge that maximally increases the score is then added (in the forward phase of the algorithm, or removed, in the backward phase). The Bayes Information Criterion (BIC) is a score that decomposes in exactly this way. For a graph G over a set of variables V and data set D over V, BIC is defined as BIC(G|D) = k ln(n) - 2 ln(L), where L is the maximum likelihood estimate of the data given the graph, k are the free parameters of the causal model and n is the number of samples. Essentially, the likelihood is penalized by a model complexity parameter k ln(n), since a complete graph (an edge between each pair of nodes) will always fit the data perfectly. For directed acyclic graphs the joint distribution P(V) over the variables V can be factorized into *P(V) =* ∏ *X*_*i*_ *in V P*(*X*_*i*_ | *pa*(*X*_*i*_)), where *pa*(*X*_*i*_) are the parents of variable *X*_*i*_ in G. As a result, BIC is decomposable into a sum of local scores of each variable given its parents: BIC(G|D) = ∑ *X*_*i*_ *in V F*(*X*_*i*_ | *pa(X_i_))* where *pa(X_i_)* are the parents of variable *X*_*i*_ in G and F is the local scoring function. If each variable is a linear function of its parents plus independent Gaussian noise, then each *P*(*X*_*i*_ | *pa(X_i_))* is a Gaussian and the local BIC score becomes 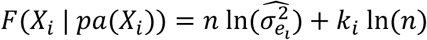 where 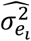 is the estimated error variance of *X*_*i*_, n is the sample size and *k*_*i*_ is the number of regressors, including the intercept. Since an added (or removed) edge changes the parent set, these *local* scores enable at each stage of the algorithm the efficient determination of the edge that maximally increases the *global* score given a current causal graph.

The fact that BIC also gives the same score to Markov equivalent graphs, and that both an edge-adding (forward) and an edge-removal (backward) phase are necessary for consistency of the FGES algorithm is not obvious, but we refer the reader to the excellent paper describing GES (Chickering, 2002) for details.

In our implementation we used the FGES algorithm published through the Tetrad code package <u>(http://www.phil.cmu.edu/tetrad/)</u> version 6.1.0. We did not force the faithfulness assumption and searched to the maximal node degree using the implemented SEMBIC score. The implementation has one free parameter *s* that functions as a sparsity parameter by multiplying the complexity term k ln(n) of the BIC score, higher values forcing sparser structures. We considered values of s from 20 down to 1, in steps of 2 for the HCP and MCP datasets, and starting from 10 down to 1 in single steps, for the smaller patient fMRI datasets.

Although the FGES algorithm does return orientation information for the causal edges in the graph, when those orientations are shared across all structures in the equivalence class, we focus in this paper principally on the adjacency structure (the causal “skeleton”). There are several reasons for this decision: (i) While (Ramsey et al., 2016) report very high accuracy (precision and recall) for the recovery of adjacency and orientation information in simulations using FGES, we show in our simulations (see **Supplementary Material**, “FGES Simulation”) that such results obtain only for the setting in Ramsey et al. (2016) using extremely sparse graphs (number of edges equal to the number of variables). We found that precision and recall measures were lower for both adjacencies and orientations, in simulations that matched the somewhat denser graphs we find here (with ~10% of possible edges present, i.e. ~500 rather than ~100 edges). See **Supplementary Figure 1** for the details of these simulation results. In particular the recall for edge orientations dropped significantly relative to the set of orientations that could theoretically be determined when switching from the extremely sparse graphs to the graph density we found here: many fewer edges that could be oriented were oriented. (ii) When we explored small subsets of real data using a (less-scalable) SAT-based causal discovery algorithm (Hyttinen et al., 2014) (see **Supplementary Material**, “SAT-based causal discovery algorithm”), we found that while the adjacency information was largely shared between FGES and the SAT-based algorithm when applied to the same dataset, the orientation information varied widely. See **Supplementary Figure 2**. (iii) Finally, the overall aim of the research plan we are outlining here is to use the electrical stimulation to provide a ground truth for orienting some of the causal adjacencies we find. In future work we hope to triangulate on the determination of causal orientation from a variety of angles. For all these reasons, we omit analyses of edge orientation in the results presented below. The entire analysis will be made available publicly, including the full FGES output.

Strategy for the discovery of reliable causal graphs. Setting a low sparsity parameter *s* in the FGES algorithm produces graphs with a larger number of edges, as one would expect, and consumes more computational time (a highly nonlinear effect). Conversely, setting a high *s* eliminates many edges but produces a graph whose edges are based on stronger evidence (vertical reliability). In essence, this is a tradeoff between sensitivity and specificity. For the HCPs dataset, we settled on a sparsity parameter *s=8,* for which ~80% of the causal graph edges were reproducible across 11 independent HCPs datasets (horizontal reliability, see below); this sparsity setting for the HCPs dataset yielded 10% of the edges of the complete graph. We also find that s=8 provides a good trade-off in the accuracy measures in our simulations on synthetic data (see **Supplementary Material**). For the MCP and patient datasets, we subsequently set sparsity settings to also produce 10% of the edges of the respective complete graphs (which corresponded to similar sparsity values for the MCP dataset, but a much lower sparsity setting for the patient dataset); see **Figure 6**.

**Figure 6.**
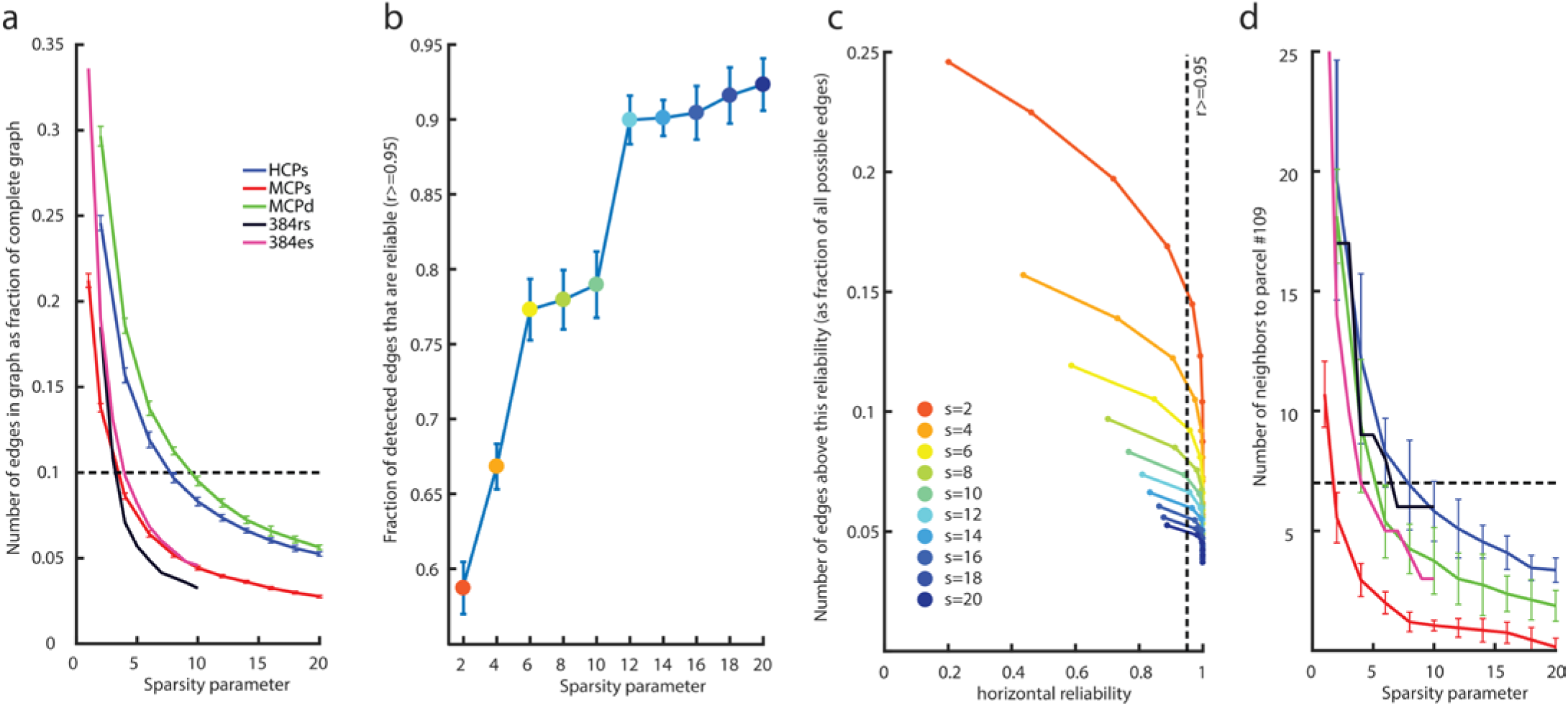
Causal graph sparsity and reliability. These panels justify how we chose the particular sparsity settings for all subsequent causal discovery analyses. a: Number of edges produced (as proportion of the complete graph) as a function of different sparsity parameters across our five datasets. As an example, the densest graph in the HCP_s_ dataset is obtained with the lowest sparsity value we tested (a sparsity of 2) and produces graphs that are about 25% complete. For our analyses in this paper, we chose a criterion of producing about 10% of the complete graph (dashed line) across all datasets as a reasonable value that permitted comparisons in all subsequent analyses. b: Fraction of edges in the HCP_s_ graph with horizontal reliability >0.95 (see **METHODS** for a description of our horizontal reliability measure). As sparsity increases (blue dots towards the right) so does the fraction of reliable edges. c: The number of edges in the HCP_s_ graph (as a proportion of the maximal number of edges that would be present in the complete graph) seen above a given horizontal reliability. Sparsity is encoded as line color. The leftmost point of each curve corresponds to observing a given edge only in 1 of the 11 datasets. Observing an edge in a sparse graph (large sparsity parameter s) is more surprising than observing an edge in a dense graph (small s), which our statistic for horizontal reliability captures (see M**ETHODS**, and **Supplementary Figure 3**): hence the leftmost 20 point of each curve shifts to the right for a higher sparsity parameter s. The second leftmost point corresponds to observing one repeat, i.e. to an edge repeating across 2 out of 11 datasets. Very high sparsity parameters (a sparsity of 20, blue curve) produce very reliable graphs. d: Same as a, except showing number of edges to the right amygdala (parcel #109), which is the node that we electrically stimulated in the neurosurgical patient. The dashed line corresponds to the number of edges to #109 obtained on average in the HCP_s_ datasets with s=8 (as in a). Equating the number of edges to #109 is another way to set the sparsity parameters across datasets, which we also explored (see **Figure 8**).

We defined horizontal reliability (in the HCPs dataset) as follows: We ran FGES on each of the 11 independent HCPs datasets for each value of the sparsity parameter. For a given sparsity value we then counted the number of times each adjacency appeared across the 11 resulting graphs, yielding for each adjacency a value from 0 to 11. We then simulated 1000 sets of 11 random graphs with the same adjacency density as the 11 real graphs of a fixed sparsity had (on average), so as to estimate how often each co-occurrence score (from 0 to 11) would occur by chance. Horizontal reliability of an adjacency A was finally defined as the proportion of adjacencies that have a lower (or equal) co-occurrence count if 11 graphs (of fixed density) were generated by chance than the co-occurrence count observed for A. See **Supplementary Figure 3** for plots of the null-distribution for different graph densities, and for the co-occurrence count that corresponds to the 95% reliability cut-off that we use in the subsequent analysis. This definition allows us to compare graphs of different sparsities more fairly, since denser graphs will necessarily present more co-occurrences by chance than sparser graphs, which is adjusted for by our estimate of the empirical chance distribution. Of course there are many ways one could define horizontal reliability measures and our measure breaks down for very dense graphs. Moreover, this measure does not capture the reliability of absences of adjacencies. Nevertheless, we found it to be a useful first pass to make adjacency reliability comparable across different graph densities for relatively sparse graphs.

Graphs with weighted edges and reconstruction of the Pearson correlation matrix. In addition to obtaining binarized adjacency matrices—graphs that either had an edge or no edge between nodes - we derived graphs that had edges with parametric weights—strong or weak causal connections corresponding to varying effect sizes. From such graphs with edge coefficients one can reconstruct the Pearson correlation matrix of the original data. The FGES algorithm returns the Markov equivalence class of causal structures, i.e. structures that all share the same (conditional) independences and dependences. Following the standard implementation in the Tetrad code package, we extracted one directed acyclic graph (DAG) from this equivalence class and then fit a maximum likelihood linear Gaussian structural equation model to the DAG using the dataset that was fed to FGES in the first place. That is, we fit a model of the form ***v* = *Bv+ e*** to the data, where v is a vector of variables representing the 110 parcels, B is a lower triangular matrix (with a zero diagonal; cf. **Figure 7**), whose non-zero entries correspond to the edges in the DAG (note that all DAGs in the equivalence class share the same adjacencies), and ***e*** is a vector of independent Gaussian errors, one for each node, with ***e** ~ N(0, **S***_e_), where ***S***_***e***_ is a diagonal covariance matrix. Essentially, the model is fit by iteratively regressing each node on the set of its parents as defined by the DAG extracted from the equivalence class.

**Figure 7.**
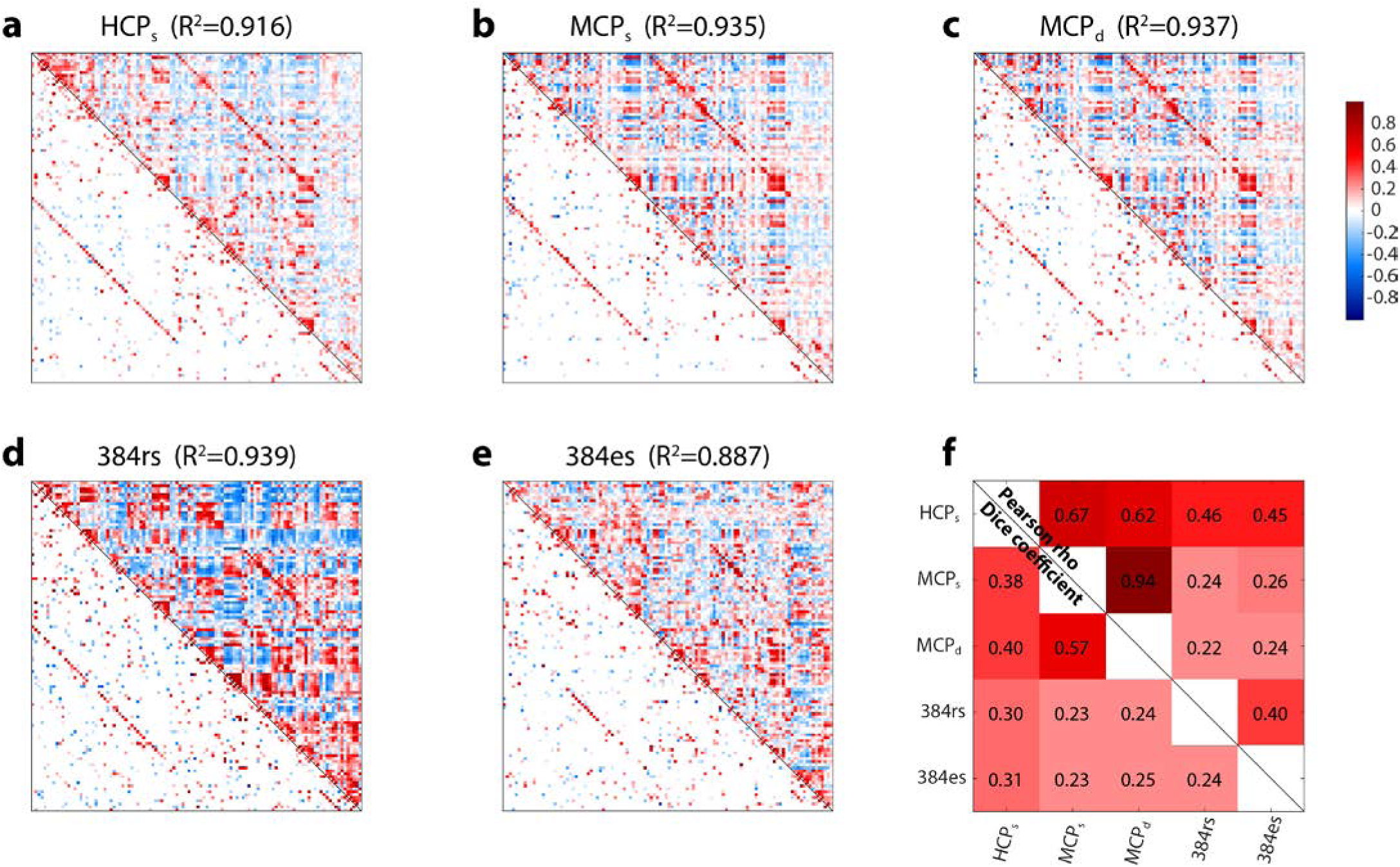
Connectivity between brain regions. a-e: Pearson correlation matrices (top triangle) and causal graphs (bottom triangle; direct connections as discovered by FGES, with weights estimated as described in the M**ETHODS** section) for all five datasets. HCP_s_: subset #1 of the sparsely sampled HCP dataset. MCP_s_: subset #1 of the sparsely sampled MCP dataset. MCP_d_: subset #1 of the densely sampled MCP dataset. 384rs: patient #384 rs-fMRI dataset. 384es: patient #384 electrical stimulation dataset. The R^2^ value at the top of each plot indicates the proportion of variance accounted for by the graph when used to reconstruct the Pearson correlation matrix (see M**ETHODS** for details). f: similarities between these 5 plots (Pearson’s r between the Fisher z-transformed correlation matrices, top triangle; proportion of edges shared in the causal graphs, bottom triangle). To calculate the proportion of shared edges in the causal graphs, we first binarized the weighted causal graphs, yielding simple adjacency matrices; for HCP and MCP datasets, we kept only reliable edges with r≥0.95 to compute overlap. We then computed the overlap across datasets using the Sorensen-Dice coefficient (***2 ∑(adjA ≠ On adjB ≠ 0)/∑(adjA ≠ 0 ∪ adjB ≠ 0***)).

We used the fully parameterized linear Gaussian structural equation model (defined by **B** and ***S_e_)*** to reconstruct the data covariance matrix ***C_v_,*** which is given by ***C_v_ = E(vv^t^***) = ***(I - B)^−1^ E(ee^t^)(I*** - *B)^−t^ = (**I*** - *B)^−1^S_e_ (I - B)^−t^* where I is the identity matrix. After standardization *C*_*v*_ can be compared to the observed Pearson Correlation matrix using the coefficient of determination *R*^*2*^ to indicate the amount of variance explained (*R*^*2*^ is shown at the top of the matrices in **Figure 7**).

Comparisons and validations. We began by making comparisons that quantify the reliability and generalizability of the methods.

(a) *Reliability of group-level whole-brain causal discovery graphs across groups of subjects in the HCP_s_ data.* We performed causal discovery separately for the 11 subsets of the HCP_s_ dataset (each derived by sparsely sampling a new group of 80 HCP subjects) and compared their graphs for different sparsity settings, as described above (**Figure 6**).

(b) *Comparison of the rs-fMRI Pearson correlation matrix to the causal graph.* The whole-brain causal graphs we produced with our criteria are relatively sparse (10% complete), and the question arises whether they indeed capture much of the structure in the Pearson correlation matrix. To address this point, we reconstructed the Pearson correlation matrix from a causal graph with weighted edges. These comparisons are shown in **Figure 7**.

(c) *HCP_s_ causal graph vs. MCP_s_ causal graph.* We wanted to see whether the graph derived from a single subject is similar to the graph derived across many subjects. We did this by comparing one sparsely sampled HCP dataset (subset #1), which comes from many (80) subjects, to an equally sparsely sampled MCP dataset (subset #1), which comes from a single subject with many (80) sessions. We kept only the horizontally reliable edges (r≥0.95) for this comparison. If the results were similar, this would confirm that the MCP dataset (from Russ Poldrack’s brain) is representative (and so can be reasonably compared to the patient’s dataset in step 2 below). This comparison is also shown in **Figure 7**.

(d) *MCP_s_ causal graph vs. MCP_d_ causal graph.* We also compared results in the sparsely sampled MCP dataset to the densely sampled MCP dataset (dense sampling being the only option with the patient’s dataset for whom much less data is available, see steps 2 & 3). This comparison addresses the question of whether autocorrelation in the data might be a problem for causal discovery, which assumes independence between samples. This comparison is also shown in **Figure 7**.

### Step 2: Causal inference from observational resting-state fMRI data in single neurosurgical patients

In this step we conducted the same analyses as in Step 1 in a new dataset: that obtained from resting-state fMRI in neurosurgical patients who required chronic invasive intracranial monitoring as part of their treatment for medically intractable epilepsy. This step was carried out in each patient who was scheduled to undergo es-fMRI (see Step 3), before electrode implantation. The motivation for this Step 2 is twofold. First, given that the patients who participate in the es-fMRI experiment are all epilepsy patients with longstanding seizures, their brain connectivity may be atypical. The rs-fMRI graph obtained from the patients can thus be used either to exclude those patients who show abnormal connectivity (or have otherwise unusual fMRI data), or as the basis for refining the graphs from Step 1 (which are all from healthy individuals without epilepsy) to better match the patients’ brain architecture. Second, the rs-fMRI data provide an independent, baseline dataset in the patients to which the es-fMRI data can be compared. An additional benefit is that the rs-fMRI data are obtained prior to electrode implantation, thus yielding signal in all parcels with a whole-brain field-of-view; by contrast, in the es-fMRI data there is typically substantial signal dropout in parcels where there are metallic contacts from depth electrodes or electrocorticography grids (including at the site of the electrical stimulation, of course).

#### Patients

We tested four neurosurgical patients who each had bilateral electrodes implanted in the amygdala. An electrocorticography (ECoG) monitoring plan was generated by the University of Iowa comprehensive epilepsy program after considering each patient’s clinical requirements. All experimental procedures were approved by the University of Iowa Institutional Review Board, who had available our gel phantom safety experiments for their evaluation prior to any human experiments (Oya et al., 2017). Written informed consent was obtained from all subjects. Patient #384 was a fully right-handed 37-year-old male; #307 a fully right-handed 29-year-old male; #303 a fully right-handed 34-year-old female; and #294 a fully right-handed 34-year-old male (see **Supplementary Table 3**). We present analysis of rs-fMRI data only from patient #384, who had the most es-fMRI runs (see below) and for whom we performed causal discovery in the es-fMRI data. We present standard GLM results of the activation evoked by es-fMRI in all four patients (Step 3; **Figure 10**).

#### Data acquisition

Resting-state fMRI runs for patient #384 were acquired on a 3 Tesla MRI scanner (Discovery 750w, GE Healthcare, Chicago, IL) with a 32 channel receive-only head coil. Each resting-state run consisted of 130 T2*-weighted EPI volumes (eyes open, central cross-hair fixated) acquired with the following parameters: TR = 2260 ms, TE = 30 ms, flip angle = 80 degrees, voxel size = 3.4 mm x 3.4 mm x 4.0 mm, 30 slices, matrix = 64 × 64, FOV = 220 mm. We obtained 5 such runs for #384. A field map (dual-echo GRE, TR = 500 ms, flip angle = 60 degrees, voxel size = 4.4 mm x 4.4 mm x 4.0 mm), high resolution T1-weighted (IR-FSPGR, TI = 450 ms, flip angle = 12 degrees, voxel size = 1.0 mm x 1.0 mm x 0.8 mm) and T2-weighted scans (CUBE TSE TR = 3200 ms, TE = maximum, echo train length = 140, voxel size = 1.0 mm isotropic) were acquired in the same session.

#### Data preprocessing and denoising

All data were minimally preprocessed using the HCP fmriVolume pipeline (v3.5.0). In summary, after rigid-body motion correction, B0 distortion correction was performed using the field map, and the mean EPI image was registered to the T1w image using boundary-based registration. All steps, including a final MNI space transformation, were concatenated and applied to the original fMRI time series in a single 3D spline interpolation step. Finally, this MNI-space time series was masked and intensity-normalized to a 4D global mean of 10000. Following this minimal preprocessing, we further applied the same denoising procedure described in Step 1 above (regression of nuisance signals and motion, bandpass filtering).

#### Causal discovery

The same analyses as in Step 1 were conducted. The major difference with Step 1 is, of course, the amount of observational data available for a single neurosurgical patient. While the HCP dataset had a large number of datapoints collected over a large number of subjects, and the MCP had a large number of datapoints collected over the course of over one year for a single subject, in the clinical setting we typically only obtained three to five 5 minute rs-fMRI runs with 130 volumes each, i.e. between 400 and 800 observations. The autocorrelation of the fMRI signal, and other factors such as high motion, further reduce the effective number of independent observations available for causal discovery. Note that we have to determine the presence or absence of 5995 (110 choose 2) adjacencies, and the edge coefficients for those that are present, and another 110 parameters for the error variances --the problem is thus underconstrained with this dataset and yields less reliable estimates (hence our proposed use of graphs derived from more data as priors, cf. **Figure 4**).

#### Comparisons and validations

Once again, we carried out a set of comparisons similar to those listed above under Step 1, but this time using the patient’s rs-fMRI dataset. We also wanted to establish that the patient has largely normal resting-state connectivity, and thus made the following comparison:

Comparison (e): HCP and MCP causal graphs vs. patient causal graph. This comparison is shown in **Figure 7**.

### Step 3: Causal inference from interventional fMRI data: es-fMRI in the amygdala

While Steps 1 and 2 relied exclusively on observational data, in Step 3 we intervene on one node of the causal network, the amygdala, using the es-fMRI technique that we recently developed. All further technical details are described in (Oya et al., 2017), and we only summarize them briefly here.

#### Patients

The same four neurosurgical patients described in Step 2. We conducted a standard whole-brain voxelwise GLM analysis of the data on all four patients (cf. below). We only carried out a (parcellated) causal graph discovery in the patient who had the most es-fMRI runs (patient #384).

#### Safety of es-fMRI

The safety of concurrent electrical stimulation and fMRI was previously established (Oya et al., 2017) through measures in a gel phantom, followed by carrying out the procedure in several patients. This demonstrated that induced currents, mechanical deflections of electrodes, and electrode or tissue heating were well controlled and all within acceptable safety levels. The electrical stimulation-fMRI experiments were performed after the final surgical treatment plan was agreed upon between the clinical team and the patient, and it was justified to move the patient to the MRI scanner (within 16 hours prior to the electrode removal surgery).

Intracranial electrodes and localization. The four patients were implanted with a combination of subdural surface strip and grid electrodes and penetrating depth electrodes; we stimulated only through the macro contacts on the depth electrodes located within the amygdala. Localization of the electrodes was done as follows. We routinely obtain two baseline (pre-implantation) structural MRI volumes, two post-electrode implantation structural MRI volumes right after implantation, another two structural MRI volumes at the time of the es-fMRI session, and a volumetric thin-sliced CT scan (1 mm slice thickness). Electrode contacts are identified on the post-implantation MRI/CT volumes and transferred onto the pre-implantation baseline MRI volumes. Great attention is paid to possible post-surgical brain shift, which is corrected with a 3D thin-plate spline warping procedure (Oya et al., 2009). For the delineation of the sub-nuclei of the amygdala, we utilized a non-linear warping applied to an atlas of the human brain (Mai et al., 1997) to draw borders of the sub-nuclei of the amygdala on the subject’s brain.

#### Electrical stimulation

Bipolar electrical stimulation was delivered through the intracranial electrodes using a battery-driven isolated constant current stimulator (IZ-2H stimulator, Tucker-Davis technology's, Alachua, FL, USA, and Model 2200 isolator, A-M systems, WA, U.S.A.). We used biphasic charge-balanced constant current stimulus waveforms of +9/-3 or +12/-4 mA, delivered at 100 Hz (5 to 9 pulses; see **Figure 5**, and **Supplementary Table 3**). Mean in-situ electrode impedance measured at the time of the experiments was 4.08 (sd = 1.65) kΩ for 100 Hz stimulation.

#### Experimental Design and data acquisition

We used a simple block design with 30s (stimulation-) ON blocks alternating with 33s (stimulation-) OFF blocks. For ON blocks, electrical stimulation was applied during a 100 ms gap between consecutive EPI volumes, when all gradients were effectively switched off; this served to minimize stimulation-induced artifacts in the fMRI data and reduce the possibility of interactions between the external electrical stimulations and RF or gradient switching-induced potentials in the electrodes. There were 10 ON blocks per run, for a total run duration of approximately 11 minutes. All scans were performed in a 3 Tesla MRI scanner using the quadrature single channel T/R head coil (patient #294: Siemens Trio; other patients: Siemens Skyra; TR = 2900 ms, TR delay = 100 ms, TE = 30 ms, flip angle = 90°, voxel size = 3.2 mm x 3.2 mm x 3.0mm, 44 slices, matrix = 68 × 68, FOV = 220 mm). During the scanning session, we carried out between one and four es-fMRI runs (#384: 4 runs; all other patients, 1 run). A T1w structural image was also acquired in the same experimental session (MP-RAGE, TR = 2530 ms, TE = 3.52 ms, TI = 100ms, flip angle = 10°, 1 mm isotropic resolution).

#### Whole-brain, voxelwise GLM analysis

We ran a standard whole-brain voxelwise GLM analysis, contrasting blocks ON and blocks OFF. Preprocessing was as described in our previous work (Oya et al., 2017). Briefly, the first two EPI volumes were discarded; slice-timing differences were compensated; motion correction was performed; retrospective denoising was applied using FIACH (Tierney et al., 2016); principal component noise regressors (n=6) were calculated and used for regressing out the effect of noise; the patient’s T1w structural volume was co-registered to that patient’s mean EPI volume; spatial smoothing with a Gaussian kernel of FWHM (full-width at half-maximum) = 8 mm was applied; EPI time series were detrended by least squares fit of Legendre polynomials of order 5; frame censoring was applied for TRs with framewise displacement >0.5 mm (Siegel et al., 2014). The hemodynamic response was modeled using a boxcar function of duration 50-90 ms (depending on the actual duration of the stimulus) convolved with a single parameter gamma function (peak at 5 s, the amplitude of the basis function was normalized to peak values of 1). These analyses were performed in subject space, with subsequent warping of the results to MNI space. Statistical parametric maps were thresholded at p<0.001(uncorrected); only clusters spanning more than 20 voxels were reported. This analysis was used in 4 patients to generate standard whole-brain, voxelwise analyses of activations evoked by amygdala stimulation (**Figure 10** below). The activation produced by the electrical stimulation showed good temporal stability across different runs within the same subject (see **Supplementary Figure 4**).

#### Parcellated analyses: causal discovery, and simple ON-OFF contrast

All es-fMRI data from patient #384 was minimally preprocessed and denoised as described in Step 1, for parcellated analyses (to match the preprocessing and denoising of the MCP dataset). BOLD signal was averaged within gray matter-masked Harvard-Oxford parcels to create parcel timeseries, and concatenated across runs. We used all collected volumes from all es-fMRI runs as samples for causal discovery analysis (cf. Step 2). We also performed a simple t-test between samples ON and samples OFF in the parcellated concatenated data, accounting for a 5s hemodynamic delay. We corrected the resulting p-values for multiple comparisons across 110 parcels using the Benjamini-Hochberg false discovery rate (FDR-corrected), and used Cohen’s d as a measure of effect size (shown in **Figure 9**).

#### Comparisons and validations

As for the comparisons we made in the preceding Step 1 and Step 2, we wanted to obtain convergent evidence, and we wanted to use the es-fMRI data to augment the causal graphs we had obtained from the prior steps. We thus compared Pearson correlation matrices as well as causal graphs at the whole-brain level to those derived from the other datasets, and we specifically examined the edges that were connected with the right amygdala across datasets. We carried out the following comparisons:

Comparison (f): All resting-state causal graphs (HCP, MCP, patient) compared with the es-fMRI causal graph. This comparison is shown in **Figure 7**.

Comparison (g): The subgraphs comprising edges connected to the right amygdala, constituting direct causal connections with other brain structures (compared across all the datasets). This is shown in **Figure 8**. These subgraphs to the amygdala do not use weighted edges and only depict whether there is an edge there or not (binary).

**Figure 8.**
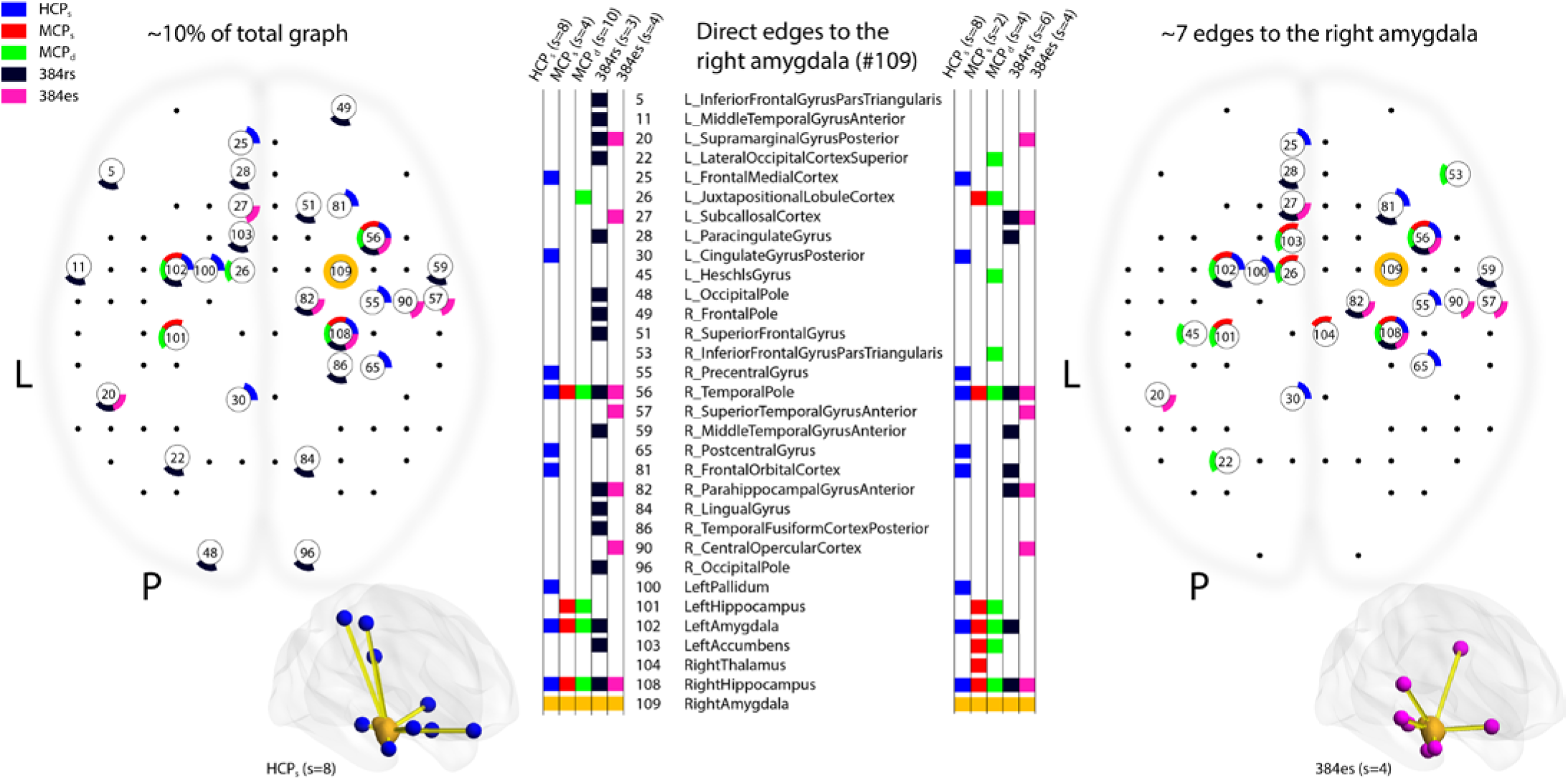
Causal discovery of direct connectivity with the right amygdala across all datasets. The two panels of this composite Figure represent the causal graph solutions obtained over our 5 datasets (5 colors, see legend at top left), showing direct edges to the right amygdala (parcel #109, circle in yellow; the same location electrically stimulated in patient #384). Left: results obtained when choosing the sparsity parameter so as to generate approximately 10% of the full graph (cf. **Figure 6**a). Right: result obtained when setting the sparsity parameter so as to generate approximately 7 direct edges to the amygdala (cf. **Figure 6**d). Sparsity settings are indicated at the top of the columns, which identify the parcels that had direct edges to the amygdala in each of the five datasets (colored entries next to the numerical and anatomical labels for all the parcels). For HCP and MCP datasets, we kept only horizontally reliable edges (r≥0.95). For the 384es dataset, we treated all brain volumes equivalently; unlike a standard contrast analysis, the causal discovery algorithm was not informed about ON and OFF states.

Comparison (h): GLM analyses of the es-fMRI compared to the causal subgraphs of edges connected to the amygdala. The GLM analyses are shown in **Figure 9** (plotted as Cohen’s d).

## Results

### Parameter setting for causal discovery and reproducibility across datasets

We first determined the reproducibility of our causal discovery analysis by deriving causal graphs from high-quality, large-sample size, statistically independent datasets. We began by using the HCPs dataset, which maximizes sample size, cross-subject generalizability, and statistical independence of the datasets. Comparing across 11 independent datasets from the HCPs, we obtained graphs at 10 different sparsity 19 settings (**Figure 6**). As expected, increasing the sparsity parameter resulted in graphs with fewer numbers of edges (moving from left to right on the x-axis of **Figure 6**a and d). As well, low sparsity resulted in graphs that were less reproducible across datasets (larger error bars at low numbers on the x-axis, **Figure 6**a and d), corresponding to increased agreement (reliability) with higher sparsity settings (**Figure 6**b). Based on these initial results, we chose to use a sparsity setting of 8 (green curve in **Figure 6**c, green marker on **Figure 6**b; see color legend inset in Figure 6c) for the HCP_s_ dataset, which yielded 10% of the edges of the complete graph (a graph with an edge between every possible pair of nodes) (dashed line in **Figure 6**a). This 10% complete graph nonetheless was able to reproduce the original Pearson correlation matrix of the dataset very well, accounting for 91.6% of the variance (**Figure 7**a). We then set the sparsity parameter in the other datasets (MCP_s_, MCP_d_, and patient; see color legend inset in Figure 6a) to a value that produced graphs with approximately this same total number of edges (10% of the complete graph, **Figure 6**a). **Figure 6** thus justifies our choice of the sparsity parameters used for the causal graphs derived from our different datasets, on which subsequent comparisons were based.

### Comparing causal graphs with Pearson correlation matrices

Before comparing causal graphs, we first compared the standard Pearson correlation (functional connectivity) matrices derived from our datasets: These comparisons are shown in **Figure 7**, as the top triangle in each of the plots. As can be seen visually in the figure’s top panel (**Figure 7**a,b and c), Pearson correlation matrices (top triangle in each plot) from our 3 large-sample resting-state datasets were very similar -we quantified this similarity using Pearson correlation as shown in the inset table (**Figure 7**f). For the patient (bottom row, top triangle in each plot), the data were considerably noisier, as expected given the much smaller number of samples, higher motion, and clinical setting. Interestingly, the patient’s rs-fMRI Pearson correlation matrix was more similar to the group-level HCP_s_ dataset (r = 0.46) than to the subject-level MCP_s_ data (r = 0.24). This was also the case for the patient’s es-fMRI whole-brain Pearson correlation matrix (r = 0.45 versus r = 0.26, see further below). We suspect that this may result from the HCP_s_ dataset smoothing out individual differences, while the MCP_s_ dataset will retain many idiosyncratic features of the one subject in that dataset (Russ Poldrack).

We next turned to the causal graphs derived by FGES for each of our datasets (bottom triangle matrices in the plots). Comparing causal graphs (bottom triangles) to Pearson correlation matrices (top triangles), visual inspection of **Figure 7** shows that the causal graph reproduces much of the structure of the correlation matrix, but is considerably sparser. Most notable at the whole-brain level are the homotopic connections, between corresponding parcels in the left and in the right hemisphere. These can be seen as the diagonal line visible in both the Pearson correlation matrix (upper triangle in each plot) and in the causal graph (lower triangle in each plot). The patient’s rs-fMRI is noisier, but again one sees this basic structure and can make out the homotopic connections. Using the Dice coefficient to quantify overlap between adjacency matrices, we found that 30% of the edges in the patient’s causal graph were also reproduced in the HCP_s_ dataset causal graph (**Figure 7**f).

### Connectivity of the amygdala

In order to provide comparisons with the electrical stimulation results, we then focused on the direct connections to the right amygdala (parcel #109) discovered by the FGES algorithm. To visualize edges connected to the right amygdala, we mapped these roughly onto a top view of the brain in **Figure 8**. Across datasets (HCPs, MCPs/MCPd, and patient #384 rs-fMRI and es-fMRI) the right amygdala was reliably found to be directly connected to the ipsilateral temporal pole and hippocampus, and the contralateral amygdala, direct connections that are supported by tracer studies in monkeys (Freese and Amaral, 2009). However, there were also numerous differences in the results from the different datasets, which will require future exploration to fully understand. One future approach we intend to incorporate is to use the causal graph inferred from one dataset (e.g., the HCP, as it may be the most reliable due to the largest number of samples provided) as a prior to help constrain the graphs obtained from other datasets (see **Figure 4**).

Finally, we examined the results of our es-fMRI experiment (**Figure 5**). Using the same parcellated data used for causal discovery (concatenated data of 4 sessions of es-fMRI in patient #384), we simply contrasted ON and OFF volumes, as typically done in a standard GLM analysis, to produce a set of node activations and deactivations. We compare these to the direct neighbors of the right amygdala found by the FGES algorithm in **Figure 9**. The results show a striking difference between the ON-OFF contrast and FGES analysis: there is no overlap at all in the sets of parcels judged by the ON-OFF contrast to be activated by the stimulation, and those judged to be causal neighbors of the stimulated amygdala by FGES. This surprising finding raises methodological questions to which we do not know the full answers yet; we discuss it further in the DISCUSSION section.

**Figure 9.**
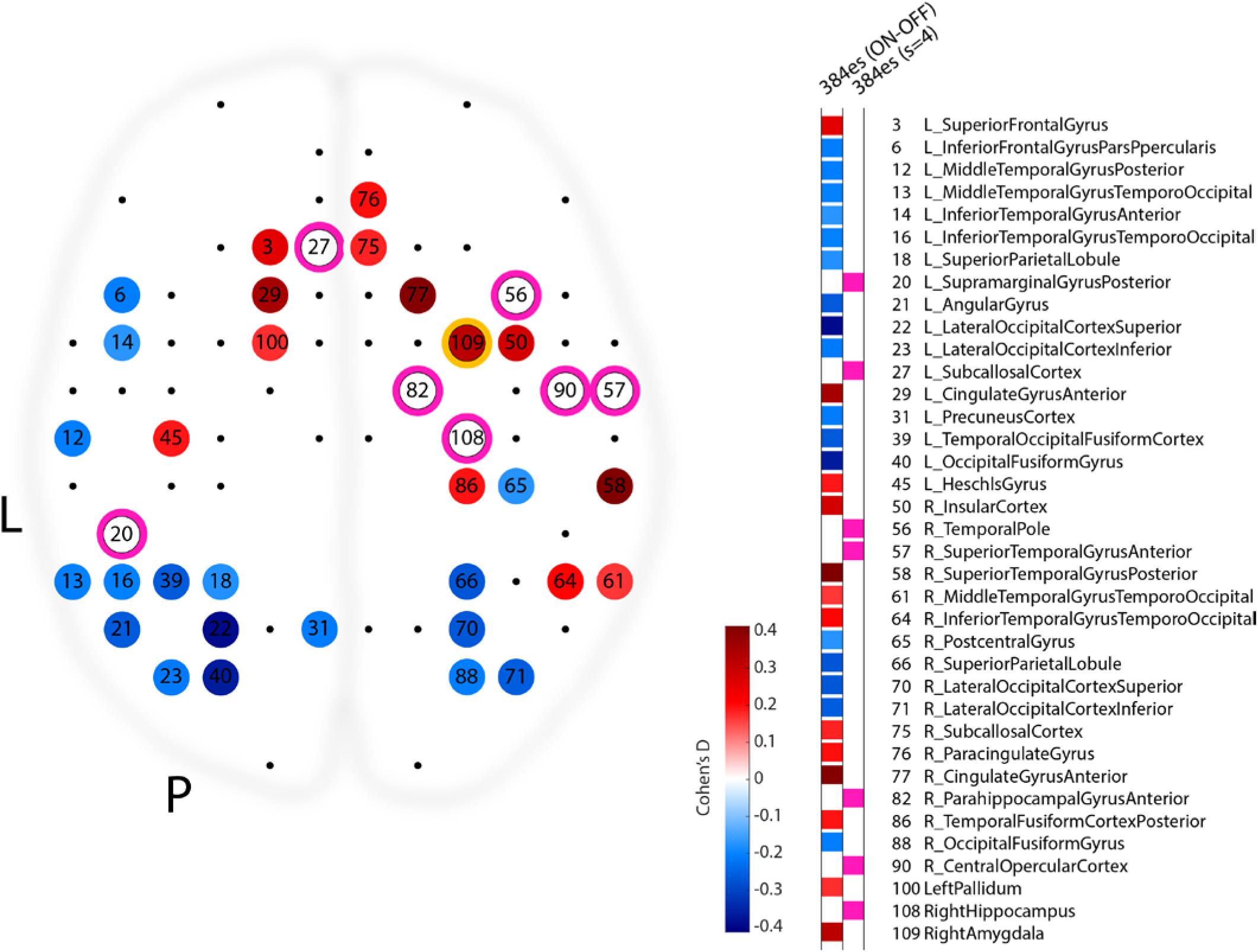
Comparison of contrast-based and causal discovery-based results for the es-fMRI data in patient #384. This view down onto a brain depicts the statistically thresholded activation produced by right amygdala stimulation (parcel #109, circled in yellow). Color of each significantly modulated node encodes the effect size of the ON-OFF contrast produced in that parcel (Cohen’s d). Nodes which are found to be directly connected to the right amygdala using the FGES algorithm are circled in pink. Unlike for the ON-OFF contrast analysis, the location and timing of the experimental manipulation (electrical stimulation of the right amygdala) did not form an explicit part of the input to the FGES algorithm as we ran it for this analysis.

To provide a more general comparison, we also show in **Figure 10** a voxelwise, whole-brain GLM analysis contrasting electrical stimulation-ON versus electrical stimulation-OFF blocks in all four neurosurgical subjects in whom we stimulated particular nuclei of the amygdala (for equivalency across patients who have differing numbers of sessions, only the first es-fMRI session was used in each patient for this **Figure 10**). Blockwise activation timecourses extracted from significantly activated ROIs showed absolute BOLD signal changes around 1% during the electrical stimulation and good reliability across runs (**Supplementary Figure 4**, and Oya et al. (2017)).

**Figure 10.**
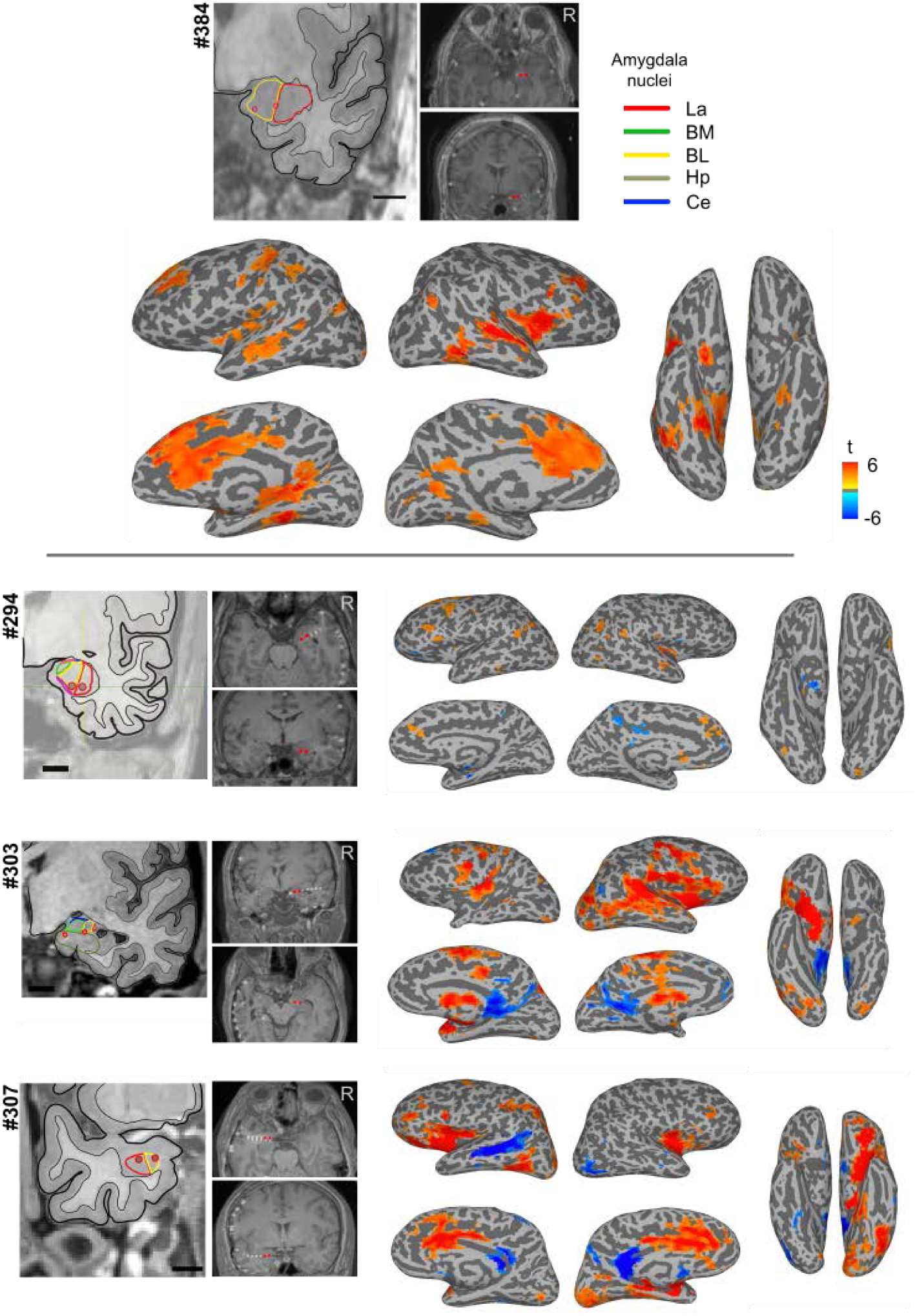
Results of concurrent electrical stimulation of amygdala nuclei and fMRI in four neurosurgical patients. Shown are GLM results from one session of es-fMRI in the four patients, as standard voxelwise, whole-brain results (mapped onto the surface). For patient #384 (top), who had four runs, we only used the first run to generate this figure. Stimulated contacts are shown as small red dots on the structural MRIs and are also shown in the leftmost panels with respect to amygdala nuclei based on a non-linear warping to a histological atlas of the human brain (Mai et al., 1997). La = lateral nucleus, BM = basomedial nucleus, BL = basolateral nucleus, Ce = central nucleus, Hp = hippocampus. Cluster-forming threshold p < 0.001 (uncorrected) with a minimum cluster size of 20 voxels. See **Supplementary** T**ABLE** 4 for the list of clusters for each patient.

Across our four patients, the results were more heterogeneous, as **Figure 10** shows. Much of this heterogeneity likely arises from differences in the specific amygdala nuclei that were stimulated. We therefore mapped the likely location of our bipolar stimulation with respect to structural MRIs of each patient’s amygdala, referenced to the Mai histological atlas (Mai et al., 1997); see *M**ETHODS*** for details. These are shown in the insets in **Figure 10**.

## Disussion

### Summary of findings

We outlined a workflow for discovering causal connections in the human brain, and provided initial validation, measures of reliability, and comparisons across datasets. We then demonstrated the application of this workflow to the connectivity of the amygdala, as a case study for the investigation of causal networks that subserve emotion processing. However, our approach is quite general and we intend it to be applicable to the investigation of any brain structure, not just the amygdala (and indeed not only to BOLD-fMRI data, or data from humans).

Our approach features two quite novel components, and suggests several further ones that were beyond the scope of the present study (cf. **Figure 4**). One novel component that can readily be applied by other researchers to sufficiently large resting-state fMRI datasets uses a causal discovery algorithm. We used a version of the fast greedy equivalence search (FGES) algorithm on rs-fMRI data parcellated into the 110 nodes obtained with the Harvard-Oxford parcellation scheme. We demonstrated excellent reliability across independent samples in two large datasets, the Human Connectome Project (HCP) dataset, and the MyConnectome dataset (MCP), and we obtained faithful reconstruction of standard Pearson correlation matrices from our sparse causal graphs (**Figure 7**).

The second component, one of the most novel aspects of our study, is the application of a new technique in human neurosurgical patients: concurrent electrical stimulation and fMRI (es-fMRI). We focused on emotion networks by investigating connectivity of the amygdala, the target of electrical stimulation. Several broad conclusions could be drawn. First, in each patient individually, there was strong consistency in the pattern of evoked BOLD activation due to amygdala stimulation: there was good session-to-session reproducibility, both in the pattern of evoked BOLD activations, and in the magnitude of the response (see **Supplementary Figure 4**, and Oya et al. (2017)). Second, there were specific differences in the statistical maps resulting from electrical stimulation across each of the four patients, as shown in **Figure 10**. This likely reflects the fact that different amygdala nuclei were stimulated in each patient (see insets in **Figure 10**), and indeed on different sides of the brain (patient #307 had left amygdala stimulation, the other three had right amygdala stimulation). However, it is also possible that there are individual differences in amygdala connectivity in the patients, a possibility especially pertinent (and clinically relevant) given that all patients had long-standing epilepsy. Studies in additional patients will be required to further understand these differences and to determine to what extent the activations seen here can be reproduced reliably across different patients in whom exactly the same amygdala nuclei are stimulated. The accrual of larger sample sizes will be required to address this issue.

The causal discovery analyses, both from resting-state data across three different datasets (HCP, MCP, and the patient #384’s pre-operative rs-fMRI), and from the rare electrical stimulation with concurrent fMRI in the four epilepsy patients, all provided novel findings about the connectivity of the amygdala. Many direct connections that would be predicted based on the known connectivity of the primate amygdala (Freese and Amaral, 2009) were also found here. For instance, there were prominent connections with temporal cortex, prefrontal cortex, and cingulate cortex. Some of the most reliable direct connections found across datasets were with the temporal pole, hippocampus, and contralateral amygdala. It is also notable that the es-fMRI results tended to produce strong activations in temporal and prefrontal cortices, but strong de-activations in posterior cingulate/retrosplenial cortices (**Figure 10**).

Perhaps most important at a global level is the finding that most of the correlations seen in standard analyses of functional connectivity (the Pearson correlation matrices shown in the top triangle plots of **Figure 7**) are due to indirect effects rather than direct causal effects of one brain region on another (cf. the much sparser causal adjacency matrix shown in the lower triangle plots in **Figure 7**). This is expected, since it is well known that not every brain region is connected to every other brain region, but that connectivity is much sparser than that.

Much more surprising was the finding that, in the es-fMRI dataset, most or all of the activations observed with standard GLM methods appear to arise from indirect causal effects, since we did not find them as direct edges in our causal discovery results (**Figure 9**, **Figure 10**). Further analyses that visualize edges that are 2- or even 3-removed from the amygdala could help to understand how the activations that we found due to electrical stimulation (**Figure 10**) arise at a network-level. It is worth noting that cortical projections from the amygdala are generally thought to be modulatory in nature: connections with temporal visual cortices terminate in superficial cortical layers (Freese and Amaral, 2006), and projections to prefrontal cortex may exert effects via the dorsomedial thalamus rather than directly (Miyashita et al., 2007). A full understanding of how network-level effects of the amygdala arise will require not only further electrical stimulation-fMRI studies, but will also require the application of other causal discovery algorithms that can incorporate feedback (see further below, and **Supplementary Material**).

Taken together, the results highlight the promise, challenge, and next steps of this novel framework. We demonstrated that causal discovery analyses can produce graphs that are reliable and that capture the correlation structure of the observational data. We also demonstrated that es-fMRI in the amygdala produces robust activations in distal brain structures. Many of the results fit with what one would expect given current knowledge of the connectivity of the amygdala: there is activation in medial prefrontal and cingulate cortices, in insula, and in temporal cortex, amongst other regions. Yet the notable differences across individual subjects also highlight the difficulty in obtaining reproducible stimulation results across patients, and in obtaining a sufficiently large number of samples for reliable causal discovery. These issues can probably be resolved partly through the accrual of more data. Other next steps consist in investigating other nodes in emotion networks, and including results from experiments in animals. We briefly comment on next steps and extensions below.

### Investigating emotions and feelings

While the present paper focuses its scope on an analysis just of data from the brain, such data will eventually need to be linked to their causal effects on the dependent measures that are typically used to infer emotions—autonomic responses, changes in facial expression, verbal reports of emotional experience, and a variety of effects on task performance (**Figure 11**). Investigating causal connections related to emotions in the brain at rest, as we did here, is clearly suboptimal, because the different nodes of the network are unlikely to be as interactive during rest as they are during emotion processing. We would thus want to apply the causal discovery methods that we document here to fMRI data that reflects brain states of putative emotions — either induced through sensory stimuli (e.g., watching emotionally laden film clips (Gross and Levenson, 1995)), volitional instruction (e.g., asking people to remember emotional autobiographical events (Damasio et al., 2000)) or through direct electrical stimulation of structures such as the amygdala (Bijanki et al., 2014; Dlouhy et al., 2015; Gloor et al., 1982; Halgren et al., 1978; Willie et al., 2016). The latter is a particularly intriguing aspect: as we demonstrated here it is in fact possible to combine electrical stimulation with concurrent fMRI measures, and it would offer the most direct test of the putative causal roles of brain structures in emotion.

**Figure 11.**
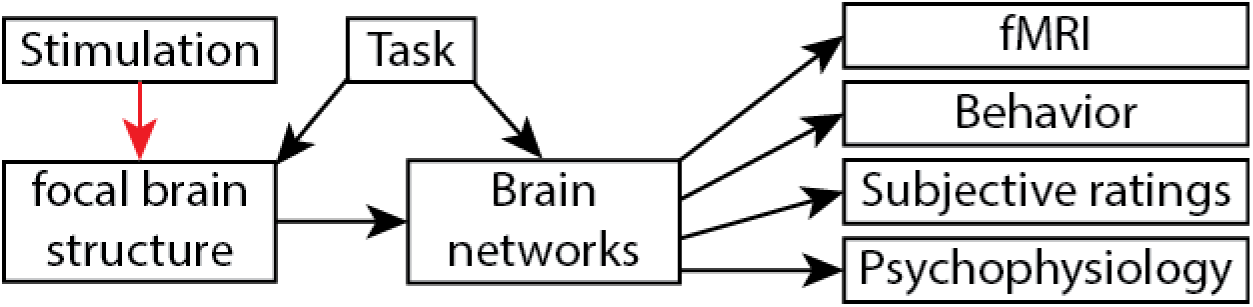
Extending the present results to a more comprehensive investigation of emotion. As also suggested in Figure 4, we would eventually want to extend the present findings to actual induction of emotion (either through direct stimulation, or suitable stimuli/tasks), and to the inclusion of other dependent measures, such as psychophysiological responses (skin-conductance response, heart-rate changes, pupillometry, etc.) and even verbal reports of the conscious experience of an emotion.

This issue would be of very high relevance to the strategic planning of deep-brain stimulation to treat mood disorders, or indeed more broadly to treat any number of severe disorders that are medically refractory and that are candidates for treatment through deep-brain stimulation. Alterations in brain connectivity are now thought to underlie much of psychopathology (Fox et al., 2014; Greicius, 2008; Greicius and Kimmel, 2012; Zhang and Raichle, 2010). Both invasive (Lozano and Lipsman, 2013) and noninvasive (Dayan and al, 2013) neurostimulation are regularly used, and gaining popularity, to treat a number of neurological and psychiatric diseases, including Parkinson’s and Alzheimer’s disease, depression (Fox et al., 2014; Mayberg et al., 2005; O'Reardon and al, 2007) and memory disorders (Hamani et al., 2008; Suthana et al., 2012). All of these avenues for treatment show a frustrating combination of features: they can be extremely effective for certain patients, yielding dramatic improvements in quality of life; but they don’t work for others, and we do not understand why. The current inadequacy in strategic planning of deep brain stimulation for treating mood disorders, and in predicting personalized outcome, stems from our ignorance of what deep brain stimulation actually does to the brain. The framework we presented here seeks to address this important outstanding question.

The approach we presented here would also help to resolve several ongoing scientific debates. For instance, the amygdala has long been thought to be necessary for fear, in humans and in animals (Amaral and Adolphs, 2016). But the evidence from lesion studies in humans does not show that the amygdala causes fear (only that absence of the amygdala interferes with it), as we noted in the INTRODUCTION.

Electrical stimulation of the amygdala, which could show that the amygdala causes fear, has been thought to act through other indirect mechanisms, for instance via stimulating white matter pathways that instead activate regions of cortex, which in turn cause the conscious experience of fear (LeDoux, 2015; LeDoux and Brown, 2017). Our es-fMRI paradigm, coupled with a causal discovery analysis, as outlined in this paper, could resolve this issue and ultimately yield an understanding of the proximal causal substrates for all the different aspects of an emotion, including its conscious experience.

Our long-term goal is to use causal discovery and es-fMRI to investigate the neural mechanisms that underlie different components of emotion. It is notable that all of the es-fMRI experiments we presented here were performed at a level of stimulation at which the patients were at chance in discriminating whether they had been stimulated or not, and no observable measures of emotion were produced (other than brain activations measured with fMRI). An experiment we plan to do next is to parametrically increase the amplitude and/or duration of the electrical stimulation. As one gradually stimulates the amygdala more and more, measurable components of emotion should be induced: there might be changes in autonomic responses such as skin-conductance response (Willie et al., 2016), changes in cognitive bias such as judgments of facial expressions (Bijanki et al., 2014), or changes in reported conscious experience (Halgren et al., 1978). What changes in the causal graph that describes the brain networks as these emotion components are induced? What accounts for the difference between stimulation trials in which the patient reports feeling nothing, and in trials in which the patient reports feeling an emotion? These are major unsolved questions in affective neuroscience that the framework we outline could begin to address.

### Limitations and assumptions

Many of the assumptions we have made in our causal analysis are unrealistic for fMRI data. The brain is known to contain many connections with feedback; it is likely that there are unmeasured confounders; it is not plausible that the actual causal connections in the brain are linear Gaussian in form; despite our comparison between sparsely sampled and densely sampled data (MCP) one may remain concerned about the i.i.d. assumption of the data. By such a standard, the present analysis can only be taken to show that rather sparse causal structures can give rise to the correlations observed resting-state fMRI data.

In the **Supplementary Material** we do explore what happens when some of these assumptions are dropped, in particular the assumption of no feedback (acyclicity) and no unmeasured confounding (causal sufficiency). We have applied a SAT-based causal discovery algorithm (Hyttinen et al., 2014) that does not make these assumptions to a small subset of three of the datasets considered in this paper (since the algorithm in its current form does not scale to large numbers of variables) and we compared the results to those from the FGES algorithm. We found that causal adjacencies are remarkably reliable when the two methods (FGES and SAT) were applied to the same dataset, but that orientations of edges in the causal graphs differed. While promising, these results also indicate that significantly more effort is needed to scale the methods with weaker background assumptions (SAT) to be applicable to datasets with many variables (such as a whole brain parcellated into ca. 100 nodes). We are actively engaged in developing more scalable versions of the SAT-based method and in the development of a fast non-parametric independence test, which would also allow us to drop the parametric assumption of linear Gaussianity. Such a test could also be used in other causal discovery algorithms that use weaker assumptions, such as the various versions of the FCI-algorithm (Spirtes et al., 2000a). These future developments will require close collaboration with experts on causal algorithm development.

There are also important limitations to note at the stimulation end. Our electrical stimulation is quite imprecise compared to alternative approaches that are possible in animal models, such as optogenetics. Although we used bipolar stimulation to constrain current spread, and although the region of activation near the site of stimulation is fairly focal, this is still a large volume of neural tissue (several cubic millimeters). Not only will this introduce imprecision in the anatomical localization of the stimulation, but it will also subsume different anatomical subdivisions and cell populations, and even fibers of passage. This issue is especially problematic in the amygdala, and may be less acute for some surface cortical sites (we are also able to stimulate through grids that are on the surface of cortex). Depending on where one is stimulating, many different circuits can be activated; or inhibitory interneurons as well as excitatory neurons can be activated; or nearby white matter can be stimulated.

Our future plans for addressing these limitations are to try to triangulate on the results with as many methods as possible. In humans, we illustrated two in the present paper. Causal discovery algorithms applied to resting-state data will still have the limitations of fMRI, but do not have the just mentioned problems associated with electrical stimulation. As such, they may be able to provide priors that can help constrain the results from electrical stimulation (cf. **Figure 4**). We are also actively exploring animal models, which will ultimately be essential to obtain sufficient resolution and control. Convergent evidence from such studies can further help with the interpretation of the results from humans. Although there are of course difficult questions about homology, it is already the case that a number of studies in nonhuman primates has given us very detailed insight into circuits related to fear and anxiety, and allowed a considerably finer fractionation both of the circuits and of the behaviors than is currently possible in humans (see (Fox and Shackman, 2017; Shackman and Fox, 2016) for an overview). The overall research program should thus incorporate results from rodents, monkeys, and humans. Each of these has somewhat complementary strengths and limitations. The rodents currently offer the most precise manipulation of circuits through optogenetics, but better methods for whole-brain imaging of activations are still needed (such as imaging using ultrasound, rather than BOLD-fMRI, for example). The monkeys are beginning also to offer optogenetic and chemogenetic approaches, although this is still more limited in application than is the case in rodents. However, monkeys are of course a better animal model for human emotions than are rodents. Finally, human studies will always be limited in the precision with which we can experimentally investigate and manipulate circuits, but offer large datasets based on fMRI and provide subjective reports of experiences—the dependent measure that also determines disorders of emotions we wish to treat.

Finally, there is no shortage of conceptual and programmatic challenges that of course need to be addressed. Transparency (ideally, pre-registration of experiments) and data sharing are two important programmatic aspects that apply to affective neuroscience as they do to all other scientific subdisciplines, and a host of challenges needs to be addressed in order to leverage fMRI results to reliable conclusions about the brains of individual people (Dubois and Adolphs, 2016). Relatedly, there is the need for more precision in the methods, the neuroanatomy, and the cognitive, behavioral, and experiential variables that can be measured. This is a very large task, but recent prescriptions in the case of the amygdala and fear give us examples of what needs to be done (Shackman and Fox, 2016).

### Comparing causal discovery with standard GLM results

One of the most striking, and unexpected, findings from our study were those shown in **Figure 9**. We found that the brain regions activated by electrical stimulation of the right amygdala in the es-fMRI dataset (as analyzed with standard GLM analysis, contrasting ON-OFF electrical stimulation) were completely nonoverlapping with the brain regions found to have direct edges (direct causal connections) with the right amygdala from FGES. Although we do not have a full explanation of this finding, it seems striking enough to warrant further investigations. We make the following remarks.

(i) The direct GLM contrast between the ON and OFF blocks depends on knowledge about which data are ON and which data are OFF, as specified by the experimenter, but is independent of the measured actual activation of the right amygdala. By contrast, FGES operates on the complement of this set of information: FGES, as we ran it, is not informed about the stimulation at all, but instead uses the measured activation of the amygdala and attempts to determine its direct causal neighbors. This is a big difference that will need to be probed further in future studies. In preliminary explorations we did run FGES with the values of the right amygdala replaced by 1/0 for ON/OFF, corresponding to the stimulation blocks (i.e. the same information available to the GLM), and we found that the right amygdala was causally disconnected from all other nodes. It is unclear what explains this result, but it is likely that the number of independent ON/OFF samples is inadequate in a blocked design (since adjacent trails are strongly depend on one another). Sparse event-related designs may circumvent some of these problems in future es-fMRI studies.

(ii) There is some overlap when FGES is run on patient #384’s es-fmri data and when it is run on the other datasets, as shown in **Figure 8**. In particular, the ipsilateral temporal pole and hippocampus are found to be connected to the amygdala across the board, a finding that is also quite consistent with what one would expect from prior studies of amygdala connectivity, including direct tracer studies in monkeys. Oddly, these two regions do not show up as significantly activated by the stimulation according to the ON-OFF GLM contrast analysis, as can be seen in **Figure 9**. Differences in equating the statistical thresholding for the GLM and the sparsity settings in our FGES analyses may also partly explain these discrepancies. Future studies should undertake a more comprehensive analysis over a larger range of thresholding and sparsity settings. As well, one could undertake a more detailed investigation that specifically probes nodes found to be directly connected in the causal analyses, and uses them as ROIs for a GLM analysis (cf. also **Figure 4**). More broadly, there are still many more comparisons required between the different sets of results, in order to gain a better understanding of which are reliable findings obtainable with all approaches, which are reliable findings but can be discovered most sensitively only with a subset of the approaches, and which are unreliable findings that show up as false positives with some approaches.

(iii) As we already noted, it is of course quite possible that many of the effects revealed with standard GLM contrasts are in fact not due to direct causal effects, but reflect indirect and possibly quite complex network-level effects. **Figure 9** shows only the direct causal neighbors of the right amygdala from the FGES analysis. Future studies could easily extend the analysis to examine nodes that are causally connected to the amygdala by one intervening node, two intervening nodes, and so forth. Of course, the larger the degrees of separation (the larger the number of intervening nodes), the larger will be the total set of nodes connected to the amygdala. In fact, it would be of interest to explore this parametrically and visualize how many degrees of separation are required before a given proportion of the complete set of nodes are connected with the amygdala (or any other structure of interest). Most broadly, such an analysis could reveal general principles of brain network architecture: what is the average degree of separation between any two places in the brain, and how are degrees of separation distributed (do they differ for cortical vs. subcortical structures)?

(iv) It would be informative to investigate directionality and reciprocity in the connections between brain regions. We largely omitted this important issue, for two reasons already noted: (1) directionality of causal effects appears much less reliable than the presence of (undirected) edges and so we omitted it for this reason, and (2) reciprocity of connections (feedback) was explicitly assumed to be absent in FGES, since that is a background assumption required by this efficient causal discovery algorithm. We relaxed both of these constraints in an exploratory analysis using a different causal discovery algorithm (SAT) in our **Supplementary Material**. However, the SAT algorithm currently does not scale to more than about 7 nodes, making it too limited for present purposes. Future development of causal discovery algorithms that are both efficient in how they scale to large numbers of variables, and that can relax background assumptions, will be essential to drive this field forward. However, even without such new algorithm development, one could extend the current studies by focusing future work on nodes revealed in the present analysis. For instance, if es-fMRI of the amygdala activates the hippocampus, then we could electrically stimulate the hippocampus to ask if we can produce a symmetrical activation in the amygdala. Such studies are certainly possible, and limited only by the density and extent of electrode coverage in the patients.

### Future Extensions

As we already noted, immediate next experiments could be analyses carried out on already collected data. The HCP and the MCP datasets can be queried in much more detail. One could investigate connectivity of other specific nodes of putative emotion networks, including not only the amygdala, but also ventromedial prefrontal cortex, insula, and other regions. One could investigate individual differences across individuals, or groups of individuals, where independent behavioral or questionnaire-based measures related to emotion processing are available (some candidates from the HCP would be the Penn emotion recognition test, and the positive affect test, which are available for this dataset). And of course one could investigate differences between neurotypical and clinical populations (e.g., HCP versus ABIDE data to compare typical healthy brain networks to those from people with autism, respectively).

A major challenge will be how to improve the reliability of causal graphs obtained from single subjects, especially patients in whom there are often a number of additional constraints. The graphs obtained from large-sample datasets such as the HCP could be used as priors to constrain the causal discovery in smaller, noisier datasets from single patients (cf. **Figure 4**). While es-fMRI will always be limited to relatively short sessions and thus small numbers of samples, one could obtain denser rs-fMRI data in the same subjects before the implantation of the electrodes, providing additional, subject-specific prior information.

It is possible to parcellate the fMRI data into a larger number of parcels (e.g., the scheme by Glasser et al. (2016) rather than the Harvard-Oxford atlas we used, an alternative and more detailed parcellation we have already explored and which is entirely feasible in large sample-size datasets). This could provide new findings that the present parcellation scheme obscured through aggregation of functionally disparate brain areas. The causal discovery algorithm we used scales relatively efficiently with sample size. It would be interesting to compare several different parcellation schemes, and to test whether more fine-grained ones essentially reproduce the coarser ones or reveal different conclusions. Ultimately, it would be extremely interesting to run FGES voxelwise over the entire brain, as also suggested in section 7 of Ramsey et al. (2016). Not only would this be informative in whether or not it reproduces results from more aggregated parcellation schemes, but it could actually be a novel source of deriving parcellations in the first place.

Finally, it is possible to stimulate not only multiple brain regions in separate sessions, but to stimulate them concurrently in a single session (or even in a specific temporal pattern). Theoretically, a relatively modest number of stimulations can very efficiently permit estimation of the causal graph (Eberhardt et al., 2005). One would like to be able to causally intervene on specific brain structures, while collecting data with the whole-brain field-of-view of fMRI (or other emerging technologies, such as ultrasound imaging), with complete freedom in the choice of brain structures. This, of course, will never be possible in human subjects, but requires the application of our approach to animal studies. The most powerful future combination will incorporate conclusions from causal discovery studies in humans with data obtained from optogenetic-fMRI in rodents (Lee et al., 2010; Liang et al., 2015). Those optogenetic-fMRI studies would also have a further large advantage over human es-fMRI, namely, the ability to stimulate genetically identified neuronal populations. Such studies could in fact explain much of the heterogeneity seen in human experiments, since these are likely to conflate many different circuits due to their anatomical and cell-type imprecision, and would substantially help us to refine emotion circuits in the brain. The complementary strengths and limitations of human and animal approaches will ultimately be required to fully map out the causal connections that underlie emotion (Adolphs and Anderson, 2018; Shackman et al., 2018).

## Author contributions

J. Dubois conducted most of the analyses and generated all data figures in the main text. H. Oya and M. Howard conducted the electrical stimulation-fMRI experiments, and H. Oya processed the resulting data. J.M. Tyszka provided help with analysis of fMRI data. F. Eberhardt conducted all causal discovery analyses. All authors discussed and planned the overall framework and analyses, and all authors jointly wrote the manuscript. The authors declare no conflict of interest.

## Acknowledgments

Supported by NIMH grant 2P50MH094258, NINDS grant 1U01NS103780-01, and a grant from the Simons Foundation Collaboration on the Global Brain to R.A.; and by NSF grant 1564330 to F.E.

## Data Sharing

The HCP and MCP datasets are publicly available. All other data and analyses are available from the authors upon request.

## Supplementary Material

### FGES Simulation

To illustrate the reliability of the FGES algorithm and to inform the choice of sparsity parameter we used in our analyses, we performed a simulation study on synthetic data that satisfied all the assumptions made by FGES. We chose variable numbers, edge densities and sample sizes to match our setting here and provide a comparison to the simulation study of (Ramsey et al., 2016). Specifically, we ran the following simulation:

- Number of variables: 110 (since that is how many ROIs we had in our parcellation)
- Number of edges per graph: {100, 200, 500} (100 edges matched the simulation of Ramsey et al 2016 where the number of edges was linear in the number of variables, 500 edges matches the graphs we found for our data, i.e. ~10% of possible edges)
- All graphs were parameterized using a linear Gaussian parameterization. We used the default settings provided for simulations in the Tetrad code package, which samples edge coefficients uniformly in (−1.5, −0.5) u (0.5, 1.5), i.e. bounded away from zero, and error variances uniformly in (1,3).
- Sample sizes: {500, 1000, 2000, 5000} (500 matches roughly the sample sizes we have for the neurosurgical patients, 2000 roughly matches the sample sizes of the MCP datasets and 5000 matches roughly the sample sizes of the HCP datasets)
- FGES sparsity parameters: {4,6,8,10,20,50} (4 matches what Ramsey et al. (2016) used, 8 is what we used here; in the plots we omit 20 and 50 for readability)

For each number of edges we sampled 10 acyclic graphs (using the Tetrad random graph generator), parameterized them, and simulated data for each sample size 5 times from each graph. We ran FGES on each dataset with each sparsity parameter. In total we thus had 3600 causal discovery runs (= 3 numEdges x 10 graphs x 4 sampleSize x 5 runs x 6 sparsityParams). In each case, FGES outputs a Markov equivalence class (MEC) of causal graphs. For each MEC, we computed

- True positive adjacencies
- False positive adjacencies
- True positive orientations (number of oriented edges FGES returns that are similarly oriented edges in the true causal graph)
- False positive orientations (number of oriented edges FGES returns for which the same to variables are not adjacent in the true graph or they are adjacent, but the edge is oriented in the other direction)
- Orientable: We determined how many of the edges in the true graph are in theory orientable using only the independence structure implied by the graph (i.e. how many edges would be orientable with infinitely many samples).

We computed adjacency precision, adjacency recall/sensitivity and adjacency specificity, as well as orientation precision and orientation recall in the way described in the inset of **Supplementary Figure 1**.

We omit orientation specificity because true negative orientations have an ambiguous value. We averaged these values for runs that had the same number of edges in the true graph and the same sample size (i.e. over 50 runs).

**Supplementary Figure 1** shows the results. We have the following remarks:

- For the extremely sparse setting (100 edges) we recover the accuracy results of Ramsey et al. (2016), but those results do not hold up for denser models.
- Orientation recall is plotted on a scale where 1 means that all edges are oriented. Of course, not all edges can be oriented even in theory. So the dashed lines indicate for the corresponding edge density, how many edges could in principle have been oriented. We see that for 100 and 200 edges, FGES attains the maximum bound, but for 500 edges it falls quite short of orienting all edges that could be oriented. Note that for 100 and 200 edges the orientation recall sometimes exceeds the maximum of the corresponding dashed line. This is due to the fact that FGES can sometimes recover from so-called violations of faithfulness.
- Sparsity parameter of s=8, which we used for our analysis of the fMRI data in the main paper, strikes us as a good compromise given these results.
- While we provide the error bars over the 50 runs for each datapoint, it should be noted that these error bars conflate variation due to (i) independently sampling multiple datasets from the same causal model, and (ii) different causal models that have the same number of edges.

**Supplementary Figure 1.**
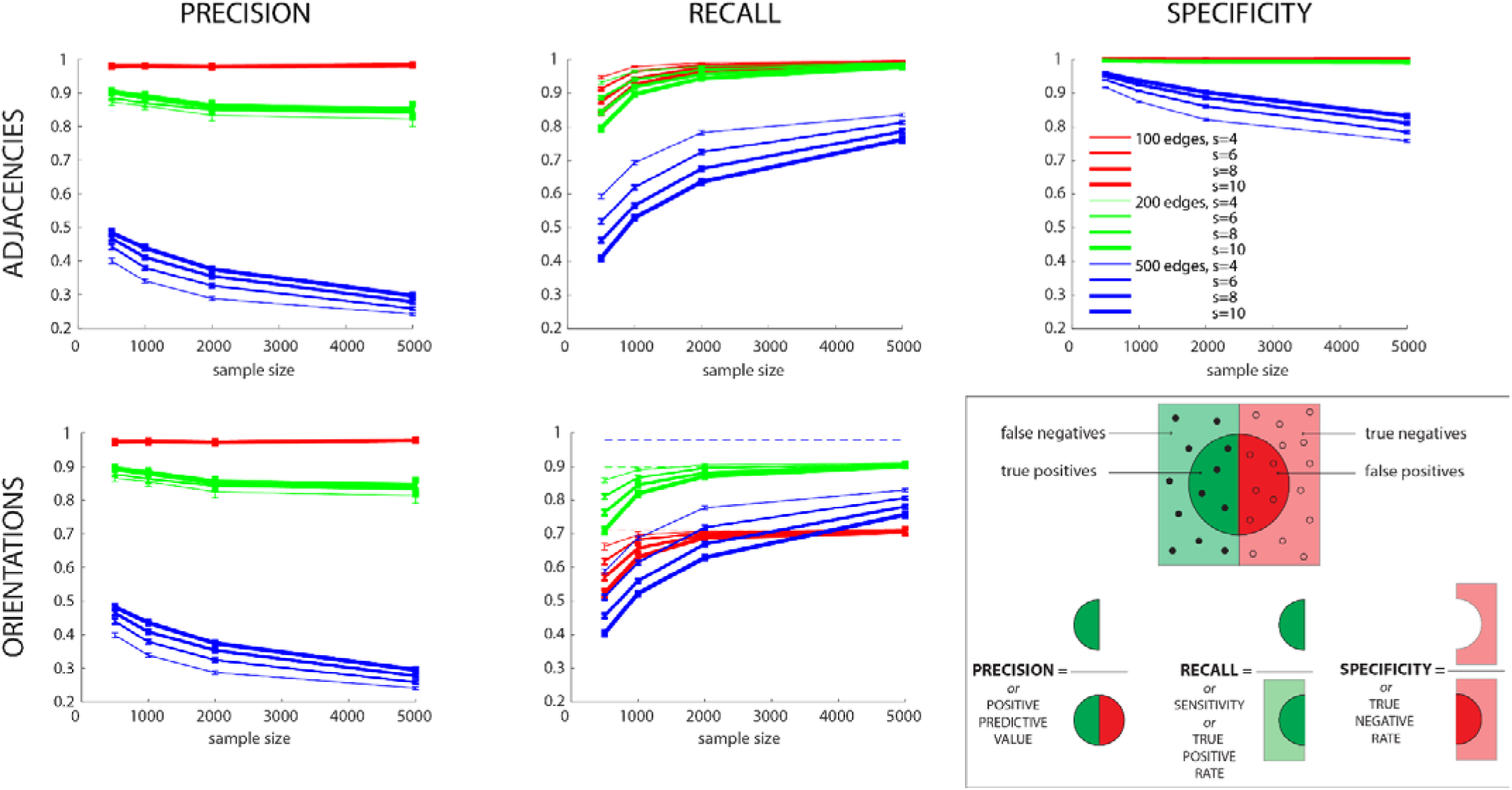
Results of our FGES simulation. We ran the FGES algorithm with sparsity parameter s=4, 6, 8 or 10, to discover a 110-node graph with 100, 200, or 500 edges. The top row shows precision, recall, and specificity (cf. inset at bottom right for a graphical reminder of what these terms mean) for adjacencies; the bottom row shows precision and recall for orientations (specificity is poorly defined for orientations). For orientation recall, the dashed lines indicate the theoretical maximum fraction of orientable edges. Error bars denote standard error of the mean (s.e.m.). See text for details.

### SAT-based causal discovery algorithm

(Hyttinen et al., 2014) developed a causal discovery algorithm based on a Boolean SAT(isfiability)-solver. Unlike FGES, it does not require the assumptions that the causal structure is acyclic (no feedback) or that it is causally sufficient (no unmeasured common causes). Moreover, it could be made completely non-parametric by using non-parametric conditional independence tests. This weakening of background assumptions comes at the cost of a significant reduction in scalability. The SAT-based method takes as input conditional independence constraints obtained from the data and translates them into Boolean constraints on the underlying graph structure. For example, a discovered independence between two variables X and Y will be translated into a constraint disallowing certain causal pathways between X and Y, while a dependence will be translated into a requirement of at least some causal connection. In general, the discovered (in)dependences may conflict due to statistical errors, so a Boolean optimization is performed to minimize the sum of (weighted) constraints not satisfied in the output causal structures. The output is a set of causal graphs, which now may also represent feedback relations and confounded variables, that are deemed an optimal fit with the data. We used the published version of the code with the most general search space (allowing for cycles and confounding), took as input all (conditional) independence tests up to conditioning set size 3, and applied a constant weight to each constraint. For the simulation here we used the implemented "classic” correlation test to determine independence.

Since the method is not scalable, we restricted ourselves to 6 ROIs that we had found to have some connection to the right amygdala (that was stimulated in patient 384), either due to the correlational analysis

(from the Pearson correlation matrices), the causal analysis (from FGES runs) or due to expert opinion (connections mentioned in the literature). Needless to say, there are more ROIs to consider, but we needed to select a manageable number.

- 109 RightAmygdala (stimulation site)
- 82 R_ParahippocampalGyrusAnterior (often correlated or causally connected)
- 56 R_TemporalPole (often correlated or causally connected)
- 108 RightHippocampus (often correlated or causally connected)
- 59 R_MiddleTemporalGyrusAnterior (often correlated or causally connected)
- 110 RightAccumbens (expert judgment)
- 81 R_FrontalOrbitalCortex (expert judgment)

For one of the HCPs datasets, for the resting state fMRI and the electrical stimulation fMRI dataset of patient 384 we ran FGES with sparsity 4 and 8, and we ran the SAT-based causal discovery algorithm allowing for the possibility of cycles and unmeasured confounding. In the case of the es-fMRI dataset we here do NOT provide the algorithms with background knowledge that variable 109 was subject to stimulation because we are not confident that the stimulation can be modeled as a surgical intervention as it is often understood in the causal modeling literature. **Supplementary Figure 2** shows the results, datasets in the rows, algorithms in the columns.

In the case of FGES the output Markov equivalence class is shown from the run when sparsity s=4. The fat edges indicate edges that were also found when s=8. Recall that all causal graphs in the same Markov equivalence class share the same adjacencies. Note that while FGES is able to orient quite a few edges in the case of the HCP dataset, it can only orient very few in the data from the surgical patient when stimulated (hence the unoriented edges).

In the case of the SAT-based algorithm, one optimal causal graph is shown, and the fat edges indicate those edges that are shared across all optimal graphs that the solver found. Orange bi-directed edges indicate the presence of an unmeasured confounder, i.e. X↔Y means that the algorithm has concluded that there is in fact some unmeasured common cause L, such that X←L→Y. Unlike in the search space that FGES considers, causal structures that are deemed equally optimal by the SAT-algorithm need not share the same adjacencies. So we also list the non-adjacencies shared by all optimal solutions, as well as the total number of optimal solutions.

We note that when comparing the two columns, there is a significant number of adjacencies that are shared, but that the SAT-algorithm resolves several of those adjacencies as due to unmeasured confounding. Note that the graphs shown for FGES result from runs only on these 7 parcels and are not the subgraphs over these 7 variables from the graph over 110 variables. So it is possible that in the graph over 110 variables, FGES would also resolve some of these adjacencies as due to common causes not included in this small subset. Of course, even a graph over 110 variables most likely does not contain all unmeasured confounders either.

Though perhaps less obvious by visual inspection, there are also a significant number of shared non-adjacencies between FGES and SAT. Importantly, these non-adjacencies imply that not only are the variables not direct causes of each other, but they are also not subject to unmeasured confounding.

Overall, we think these results vindicate our focus on the adjacency structure that FGES supplies, but we are also excited by the possibilities that lie ahead in more thoroughly exploring the data we have with more general causal discovery algorithms.

**Supplementary Figure 2.**
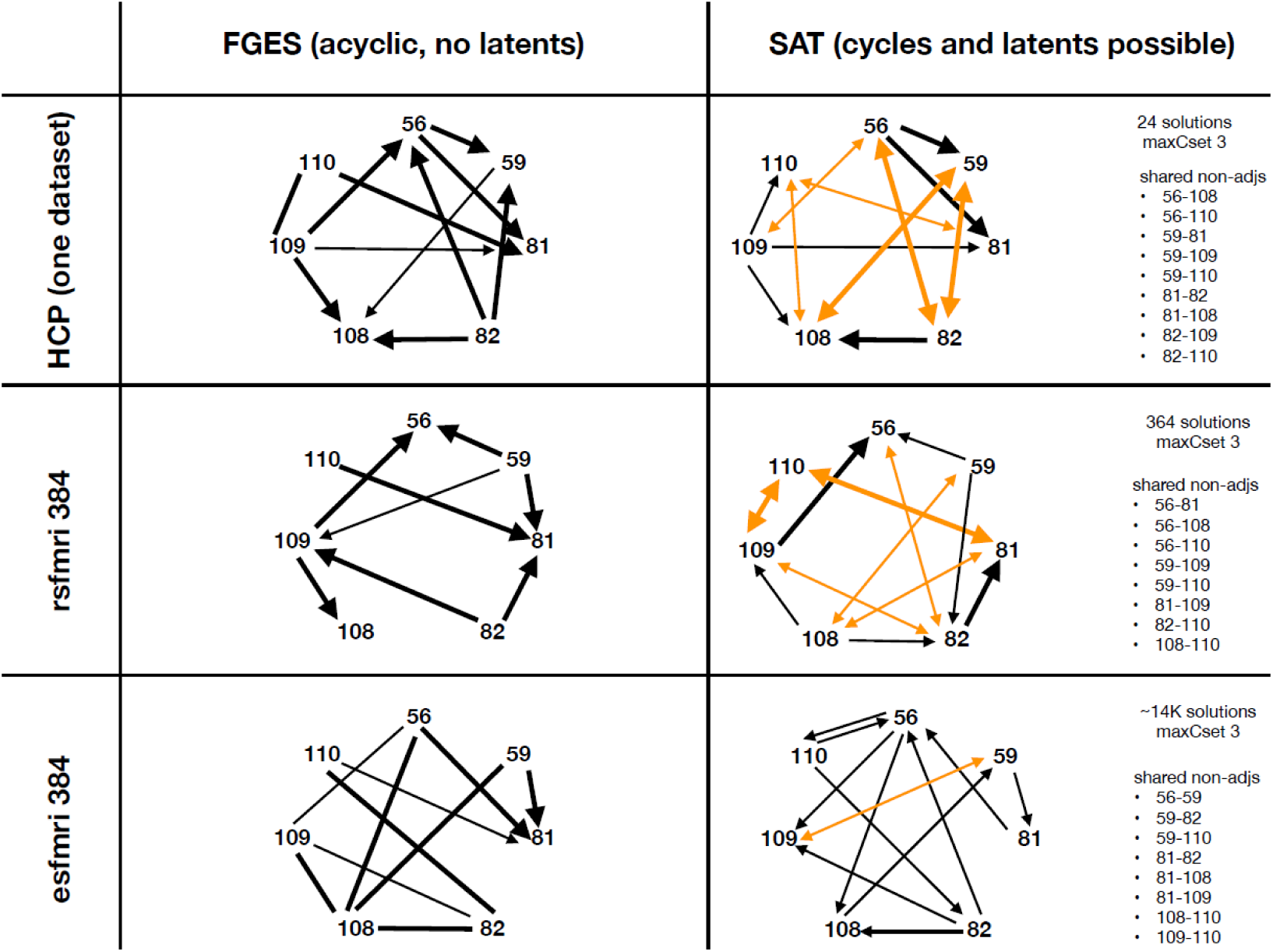
Comparison of FGES and SAT-based causal discovery algorithms, using a small set of parcels including the right amygdala (109), which was stimulated in patient #384. See text for details.

### Horizontal reliability metric

We defined horizontal reliability (in the HCP_s_ dataset) as follows: We ran FGES on each of the 11 independent HCP_s_ datasets for each value of the sparsity parameter. For a given sparsity value we then counted the number of times each adjacency appeared across the 11 resulting graphs, yielding for each adjacency a value from 0 to 11. We then simulated 1000 sets of 11 random graphs with the same adjacency density as the 11 real graphs of a fixed sparsity had (on average), so as to estimate how often each co-ocurrence score (from 0 to 11) would occur by chance (blue histograms, **Supplementary Figure 3**). Horizontal reliability of an adjacency A was finally defined as the proportion of adjacencies that have a lower (or equal) co-occurrence count if 11 graphs (of fixed density) were generated by chance than the co-occurrence count observed for A (orange curves, **Supplementary Figure 3**). This definition allows us to compare graphs of different sparsities more fairly, since denser graphs will necessarily present more co-occurrences by chance than sparser graphs, which is adjusted for by our estimate of the empirical chance distribution (see examples of a 5%, 10%, and 30% complete graph in **Supplementary Figure 3**). Of course there are many ways one could define horizontal reliability measures and this measure breaks down for very dense graphs. Moreover, this measure does not capture the reliability of absences of adjacencies. Nevertheless, we found it to be a useful first pass to make adjacency reliability comparable across different graph densities for relatively sparse graphs.

**Supplementary Figure 3.**
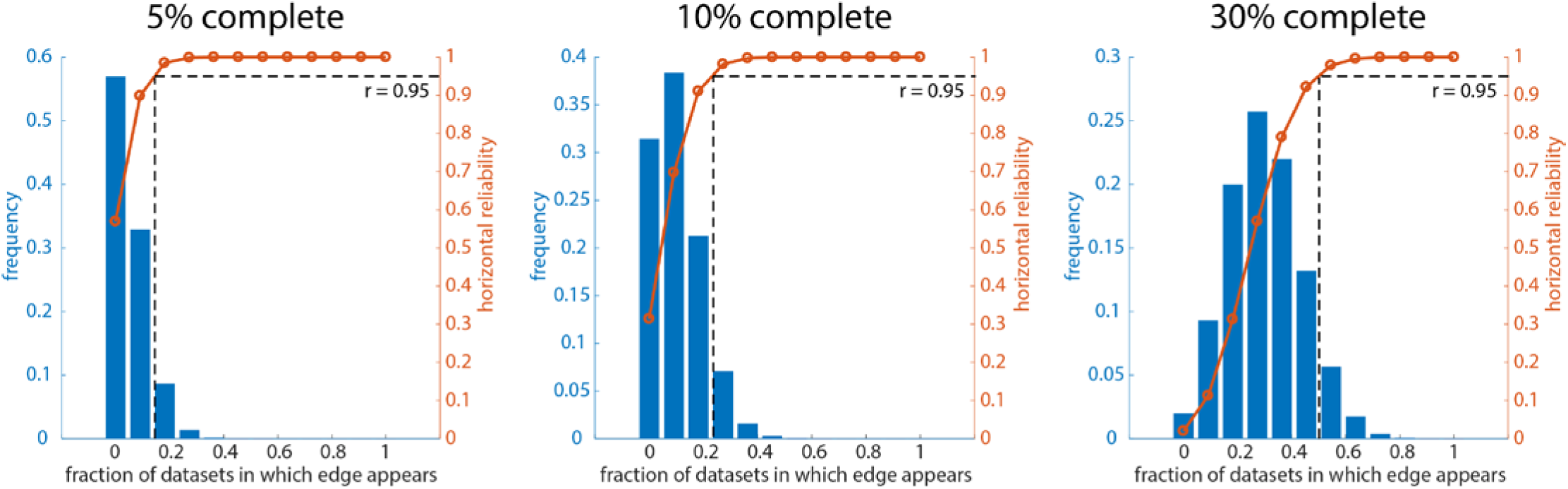
A horizontal reliability metric that can be compared across graphs of different densities. We simulated 1000 sets of 11 random graphs with three adjacency densities: 5%, 10% and 30%, from left to right. The blue histogram shows the distribution of co-occurrence scores (how many times an adjacency is repeated in the 11 subsets, from 0 to 11) that would occur by chance. The histograms differ for graphs of different densities (e.g., compare leftmost and rightmost plots). To account for this, we defined “horizontal reliability” of an adjacency A as the proportion of adjacencies that have a lower (or equal) co-occurrence count if 11 graphs (of fixed density) were generated by chance than the co-occurrence count observed for A (orange curves). Reliable edges are defined as edges with a horizontal reliability of .95 or higher (horizontal dashed lines), which corresponds to a different actual count of repeats for each graph density (vertical dashed lines).

### Reliability of es-fMRI results

We examined the reliability of the es-fMRI results within each patient across runs, in order to estimate the temporal stability of our results. At least within the timeframe of the experiment, we obtained good reliability, suggesting that the results obtained are not unstable. Future work will be required to extend this question to longer time periods, and to explore how activation maps, and connectivity, might change as a function of states such as attention and emotion.

**Supplementary Figure 4.**
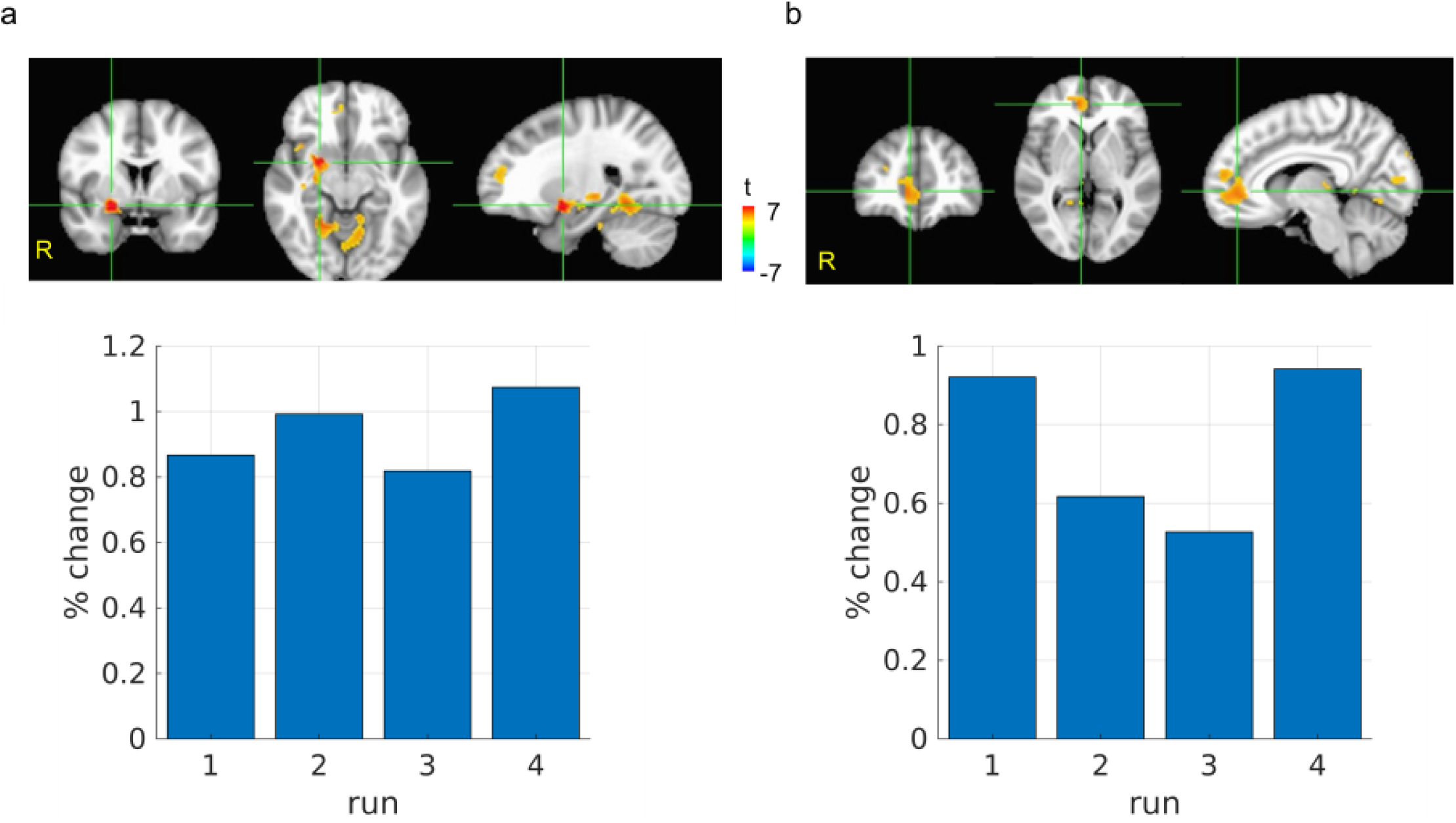
Reliability of es-fMRI results across runs. Shown are the evoked mean percent signal change observed across four runs in patient #384, at two brain locations. a: Reliability at a voxel near the stimulation site. b: Reliability at a voxel at a distal location activated by electrical stimulation.

**Table 1.**
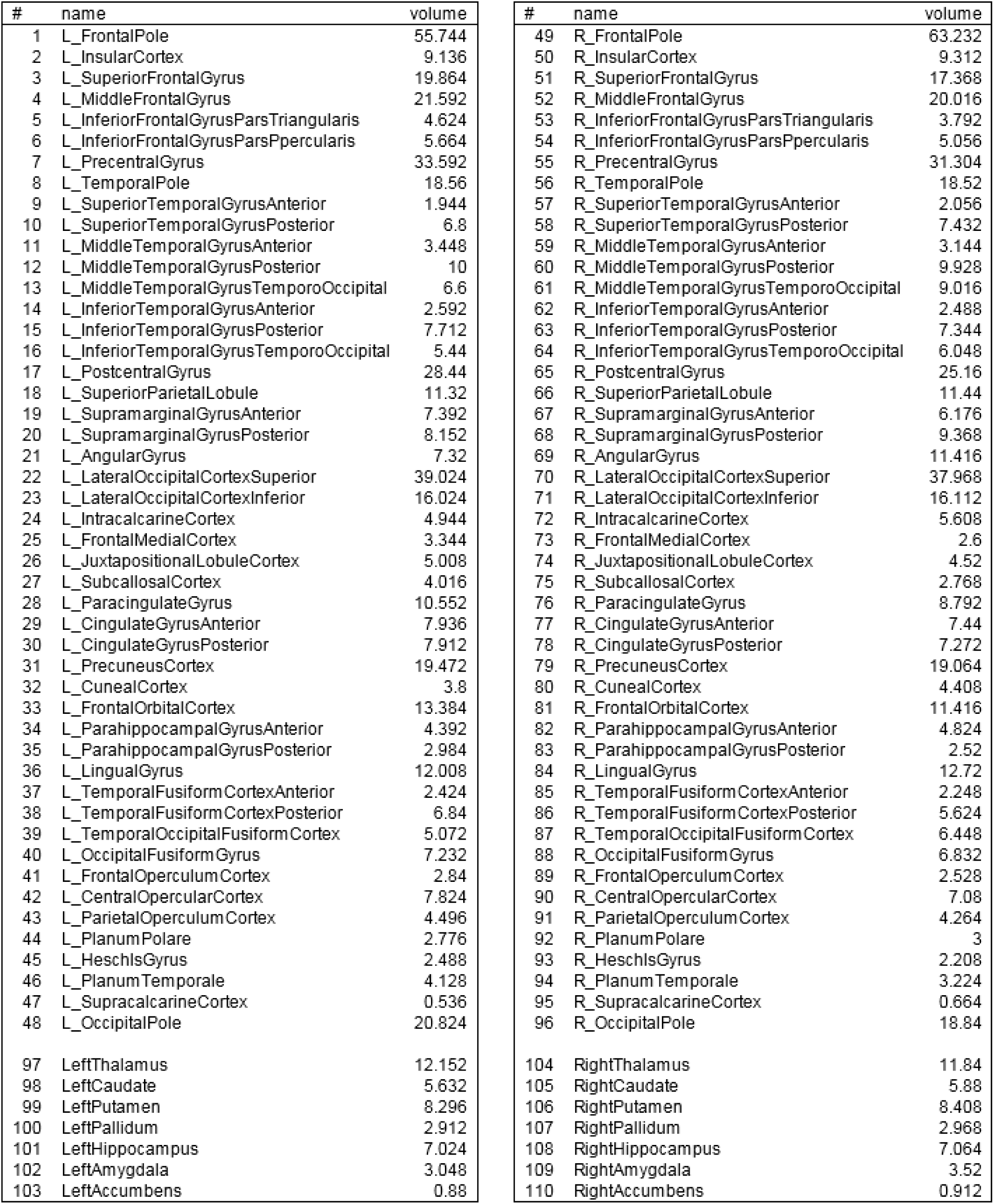
List of parcels used in this paper (from Harvard-Oxford anatomical parcellation). Volume is in cubic centimeters.

**Table 2.**
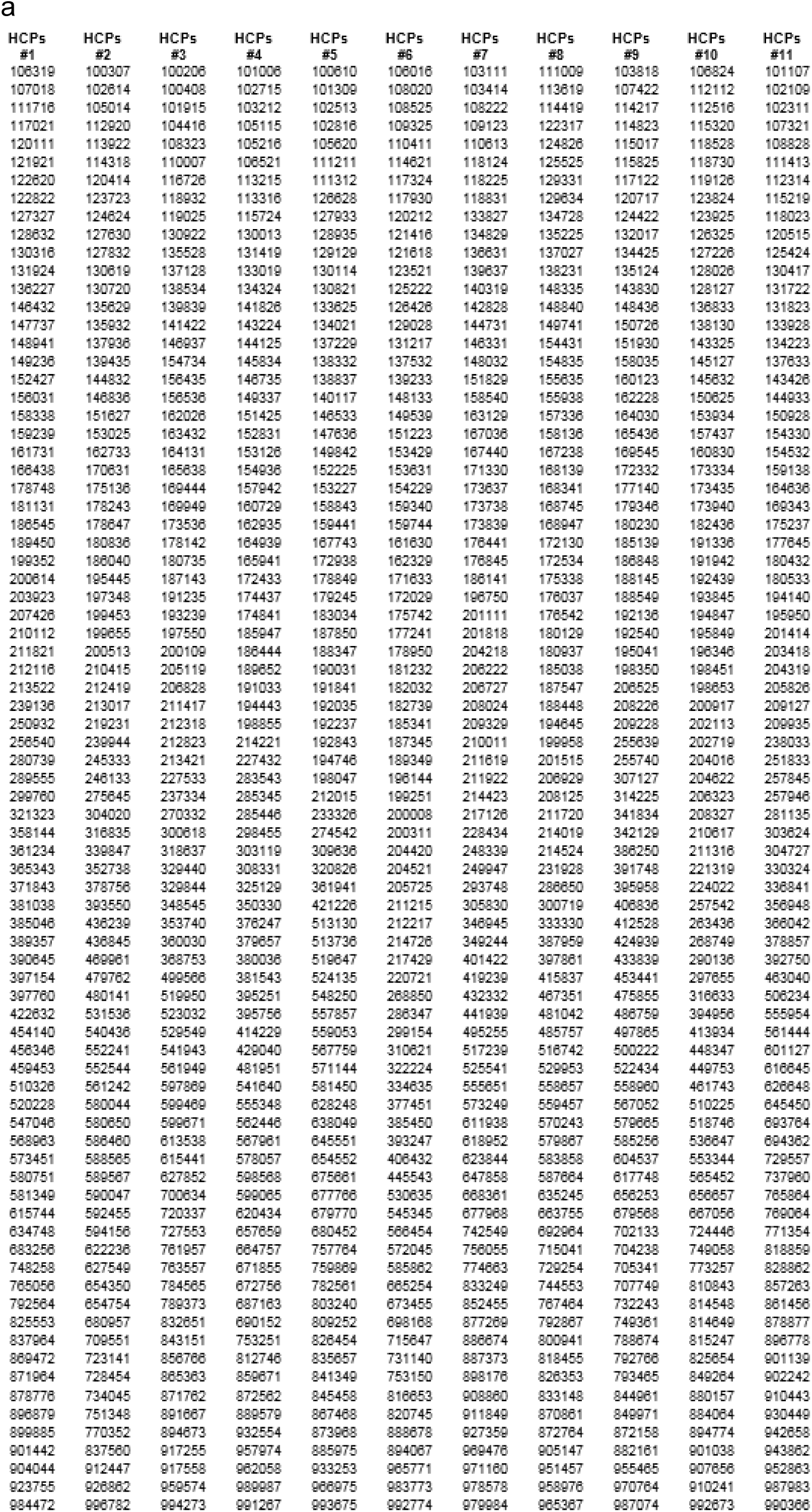
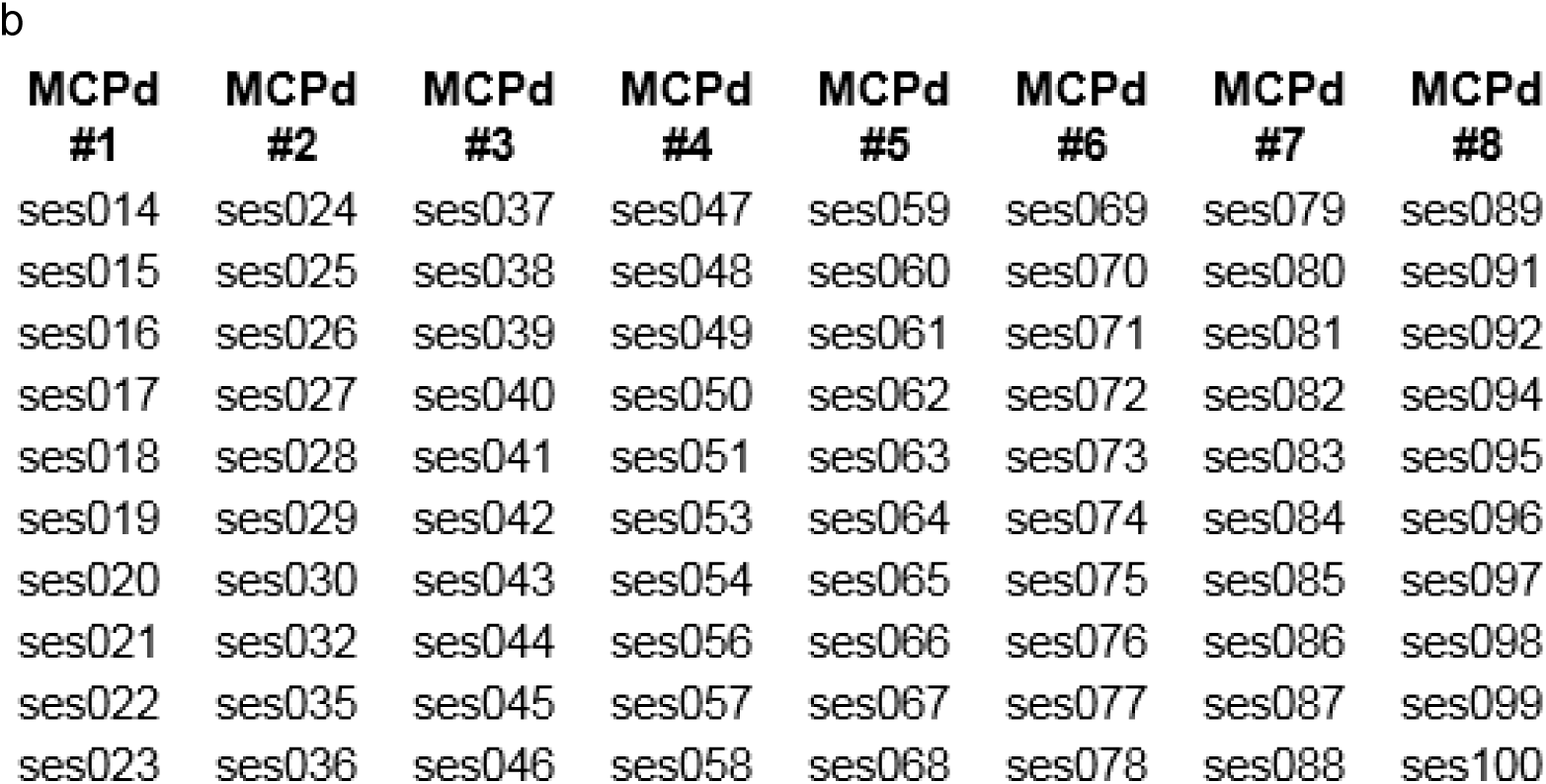
Detailed identifiers for large resting-state datasets used. a: List of 880 HCP subjects used for Dataset #1 (HCP_s_), broken down by the 11 subsets we used for our “horizontal” reliability analysis. For each subject, rfMRI_REST1_RL and rfMRI_REST2_RL runs were used, volumes [35:35:1190] b: List of 80 sessions from the MyConnectome project that we used for Datasets #2 and #3 (MCP_s_ and MCP_d_).

**Table 3.**
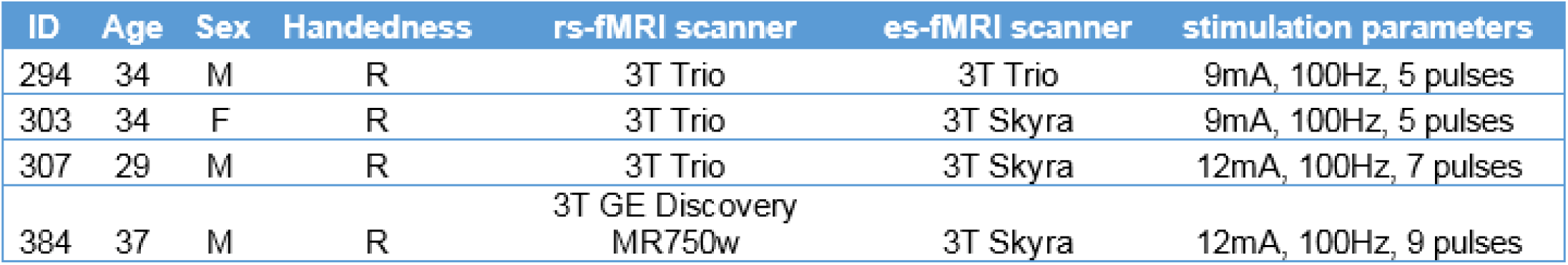
Demographics and stimulation parameters for patients included in es-fMRI study. All patients were stimulated in the amygdala.

**Table 4.**
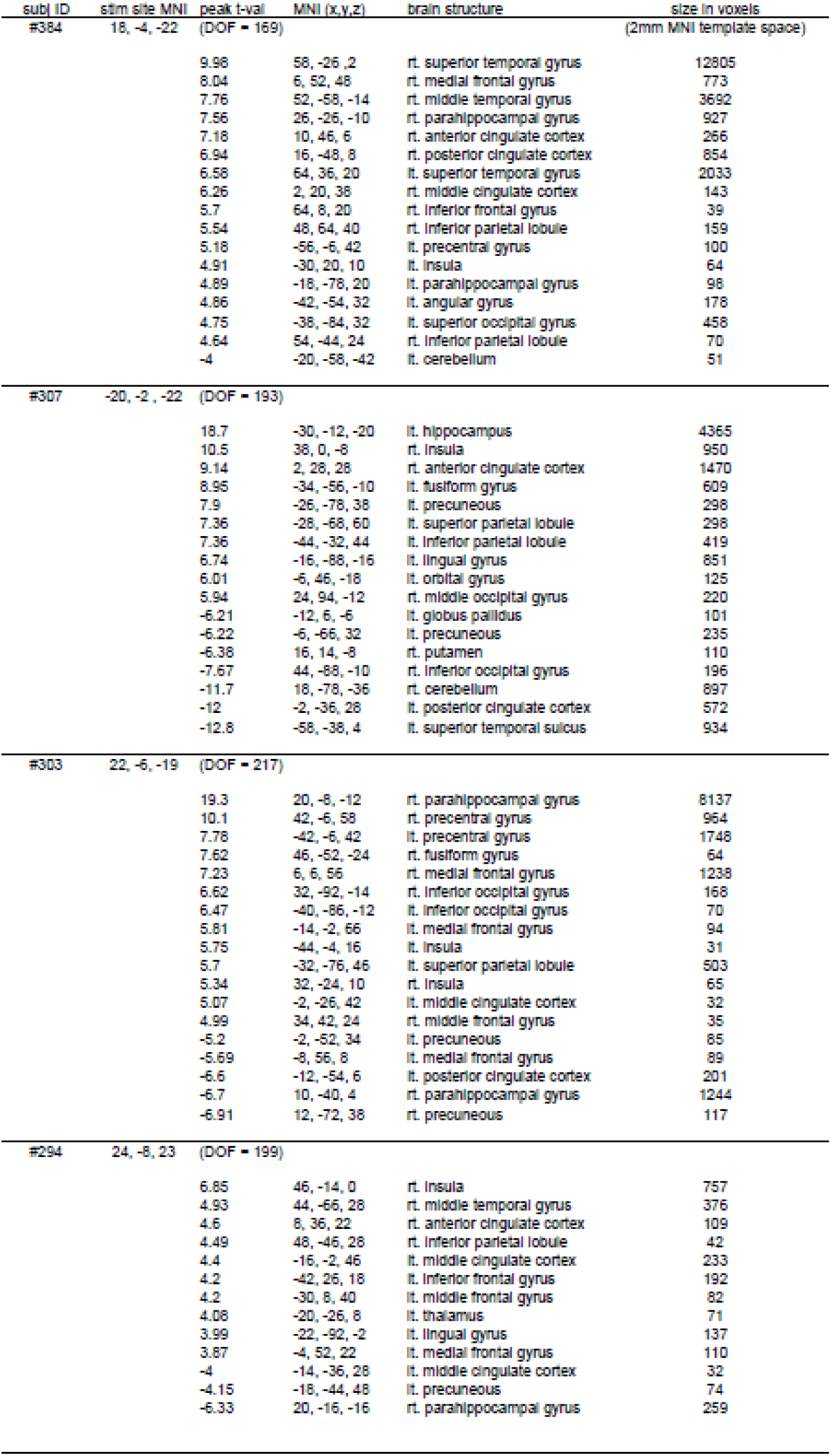
List of clusters activated by electrical stimulation (see **Figure 10** in main text).

## References

Adolphs, R. (2017a). How should neuroscience study emotions? By distinguishing emotion states, concepts, and experiences. Social Cognitive and Affective Neuroscience 12, 24–31.

Adolphs, R. (2017b). Reply to Barrett: affective neuroscience needs objective criteria for emotions. Social Cognitive and Affective Neuroscience 12, 32–33.

Adolphs, R., and Anderson, D.J. (2018). The Neurobiology of Emotion: A New Synthesis (Princeton, NJ: Princeton University Press).

Adolphs, R., Tranel, D., Damasio, H., and Damasio, A. (1994). Impaired recognition of emotion in facial expressions following bilateral damage to the human amygdala. Nature 372, 669–672.

Amaral, D.G., and Adolphs, R., eds. (2016). Living without an amygdala (New York: Guilford Press).

Barrett, L.F. (2017a). Functionalism cannot save the classical view of emotion. Social Cognitive and Affective Neuroscience 12, 34–36.

Barrett, L.F. (2017b). The theory of constructed emotion: an active inference account of interoception and categorization. Social Cognitive and Affective Neuroscience 12, 17–23.

Bechara, A., Tranel, D., Damasio, H., and Damasio, A.R. (1996). Failure to respond autonomically to anticipated future outcomes following damage to prefrontal cortex. Cerebral Cortex 6, 215–225.

Bijanki, K.R., Kovach, C.K., McCormick, L.M., Kawasaki, H., Dlouhy, B.J., Feinstein, J., Jones, R.D., and Howard, M.A. (2014). Case report: stimulation of the right amygdala induces transient changes in affective bias. Brain Stimulation 7, 690–693.

Chickering, D.M. (2002). Optimal structure identification with greedy search. Journal of Machine Learning Research 3, 507–554.

Cox, R. (2012). AFNI: what a long strange trip it's been. Neuroimage 62, 743–747.

Damasio, A., Grabowski, T., and al., e. (2000). Feeling emotions: subcortical and cortical brain activity during the experience of self-generated emotions. Nature Neuroscience 3, 1049–1056.

Davis, M. (1992). The role of the amygdala in fear and anxiety. Annu Rev Neurosci 15, 353–375.

Dayan, E., and al, e. (2013). Noninvasive brain stimulation: from physiology to network dynamics and back. Nature Neuroscience 16, 838–844.

Dlouhy, B.J., Gehlbach, B.K., Kreple, C.J., Kawasaki, H., Oya, H., Buzza, C., Granner, M.A., Welsh, M.J., Howard, M.A., Wemmie, J.A., et al. (2015). Breathing inhibited when seizures spread to the amygdala and upon amygdala stimulation. The Journal of Neuroscience 35, 10281–10289.

Dubois, J., and Adolphs, R. (2016). Building a science of individual differences from fMRI. Trends in Cognitive Sciences 20, 425–443.

Eberhardt, F. (2017). Introduction to the foundations of causal discovery. International Journal of Data Science and Analytics 3, 81–91.

Eberhardt, F., Glymour, C., and Scheines, R. (2005). On the number of experiments sufficient and in the worst case necessary to identify all causal relations among N variables. In Proceedings of the 21st Conference on Uncertainty in Artificial Intelligence (UAI), F. Bacchus, and T. Jaakkola, eds. (Arlington, VA: AUAI Press), pp. 178–184.

Feinstein, J.S., Adolphs, R., Damasio, A., and Tranel, D. (2011). The human amygdala and the induction and experience of fear. Current Biology 21, 34–38.

Feinstein, J.S., Buzza, C., Hurlemann, R., Follmer, R.L., Dahdaleh, N.S., Coryell, W.H., Welsh, M.J., Tranel, D., and Wemmie, J.A. (2013). Fear and panic in humans with bilateral amygdala damage. Nat Neurosci 16, 270–272.

Fisher, R.A. (1990). Statistical Methods: Experimental Design and Scientific Inference (New York: Oxford University Press).

Fox, A.S., and Shackman, A.J. (2017). The central extended amygdala in fear and anxiety: closing the gap between mechanistic and neuroimaging research. Neuroscience Letters in press.

Fox, M.D., Buckner, R.L., Liu, H., Chakravarty, M.M., Lozano, A.M., and Pascual-Leone, A. (2014). Resting-state networks link invasive and noninvasive brain stimulation across diverse psychiatric and neurological diseases. PNAS E4367–E4375.

Frässle, S., and al, e. (2015). Test-retest reliability of dynamic causal modeling for fMRI. Neuroimage 117, 56–66.

Frässle, S., and al., e. (2017). Regression DCM for fMRI. Neuroimage 155, 406–421.

Freese, J.L., and Amaral, D.G. (2006). Synaptic organization of projections from the amygdala to visual cortical areas TE in the Macaque monkey. The Journal of comparative neurology 496, 655–667.

Freese, J.L., and Amaral, D.G. (2009). Neuroanatomy of the primate amygdala. In Neuroanatomy of the primate amygdala, P.J. Whalen, and E.A. Phelps, eds. (New York: Guilford Press).

Friston, K.L. (2011). Functional and effective connectivity: a review. Brain Connectivity 1, 13–36.

Friston, K.L., Moran, R., and Seth, A.K. (2013). Analysing connectivity with Granger causality and dynamic causal modelling. Current Opinion in Neurobiology 23, 172–178.

Glasser, M., Coalson, T., Robinson, E., Hacker, C., Harwell, J., Yacoub, E., Ugurbil, K., Andersson, J., Beckmann, C., Jenkinson, M., et al. (2016). A multi-modal parcellation of human cerebral cortex. Nature 536, 171–178.

Glasser, M.F., Sotiropoulos, S.N., Wilson, J.A., Coalson, T.S., Fishl, B., Andersson, J.L., and Jenkinson, M. (2013). The minimal preprocessing pipelines for the Human Connectome Project. Neuroimage 80, 105–124.

Gloor, P., Olivier, A., Quesney, L.F., Andermann, F., and Horowitz, S. (1982). The role of the limbic system in experiential phenomena of temporal lobe epilepsy. Ann Neurol 12, 129–144.

Gordon, E.M., Laumann, T.O., Adeyemo, B., Huckins, J.F., Kelley, W.M., and Petersen, S.E. (2014). Generation and evaluation of a cortical area parcellation from resting-state correlations. Cerebral Cortex advanced access October 14, 2014.

Granger, C.W.J. (1969). Investigating causal relations by econometric models and cross-spectral methods. Econometrica.

Grayson, D.S., Bliss-Moreau, E., Machado, C.J., Bennett, J., Shen, K., Grant, K.A., Fair, D.A., and Amaral, D.G. (2016). The Rhesus monkey connectome predicts disrupted functional networks resulting from pharmacogenetic inactivation of the amygdala. Neuron 91, 453–466.

Greicius, M.D. (2008). Resting-state functional connectivity in neuropsychiatric disorders. Current Opinion in Neurology 21, 424–430.

Greicius, M.D., and Kimmel, D.L. (2012). Neuroimaging insights into network-based neurodegeneration. Current Opinion in Neurology 25, 727–734.

Gross, J.J., and Levenson, R.W. (1995). Emotion elicitation using films. Cognition and Emotion 9, 87–107.

Halgren, E., Walter, R.D., Cherlow, D.G., and Crandall, P.H. (1978). Mental phenomena evoked by electrical stimulation of the human hippocampal formation and amygdala. Brain 101, 83–117.

Hamani, C., McAndrews, M.P., Cohn, M., Oh, M., Zumsteg, D., Shapiro, C.M., Wennberg, R.A., and Lozano, A.M. (2008). Memory enhancement induced by hypothalamic/fornix deep brain stimulation. Annals of Neurology 63, 119–123.

Hyttinen, A., Eberhardt, F., and Järvisalo, M. (2014). Constraint-based Causal Discovery: Conflict Resolution with Answer Set Programming. Paper presented at: 30th Conference on Uncertainty in Artificial Intelligence (UAI).

Kennedy, S.H., and al, e. (2011). Deep brain stimulation for treatment-resistant depression: follow-up after 3 to 6 years. American Journal of Psychiatry 168, 502–510.

Koenigs, M., and Tranel, D. (2007). Irrational economic decision-making after ventromedial prefrontal damage: evidence from the ultimatum game. The Journal of Neuroscience 27, 951–956.

Koenigs, M., Young, L., Adolphs, R., Tranel, D., Cushman, F., Hauser, M., and Damasio, A. (2007). Damage to the prefrontal cortex increases utilitarian moral judgments. Nature 446, 908–911.

Kragel, P.A., and LaBar, K.S. (2015). Multivariate neural biomarkers of emotional states are categorically distinct. Social Cognitive and Affective Neuroscience 10, 1437–1448.

Laumann, T.O. (2015). Functional System and Areal Organization of a Highly Sampled Individual Human Brain. Neuron 87, 657–670.

LeDoux, J. (2015). Anxious (New York: Viking (Penguin/Random House)).

LeDoux, J., and Brown, R. (2017). A higher-order theory of emotional consciousness. PNAS, E2016–E2025.

Lee, J.H., Durand, R., Gradinaru, V., Zhang, F., Goshen, I., Kim, D.-S., Fenno, L.E., Ramakrishnan, C., and Deisseroth, K. (2010). Global and local fMRI signals driven by neurons defined optogenetically by type and wiring. Nature 465, 788–792.

Liang, Z., Watson, G.D.R., Alloway, K.D., Lee, G., Neuberger, T., and Zhang, N. (2015). Mapping the functional network of medial prefrontal cortex by combining optogenetics and fMRI in awake rats. Neuroimage 117, 114–123.

Lin, D., Boyle, M.P., Dollar, P., Lee, H., Lein, E.S., Perona, P., and Anderson, D.J. (2011). Functional identification of an aggression locus in the mouse hypothalamus. Nature 470, 221–226.

Lindquist, K.A., Wager, T.D., Kober, H., Bliss-Moreau, E., and Feldman Barrett, L. (2012). The brain basis of emotion: a meta-analytic review. Behavioral and Brain Sciences 35, 121–143.

Lozano, A.M., and Lipsman, N. (2013). Probing and regulating dysfunctional circuits using deep brain stimulation. Neuron 77, 406–424.

Mai, J.K., Assheuer, J., and Paxinos, G. (1997). Atlas of the human brain (San Diego: Academic Press).

Mayberg, H.S., Lozano, A.M., Voon, V., McNeely, H.E., Seminowicz, D., Hamani, C., Schwalb, J.M., and Kennedy, S.H. (2005). Deep brain stimulation for treatment-resistant depression. Neuron 45, 651–660.

Miyashita, T., Ichinohe, N., and Rockland, K.S. (2007). Differential modes of termination of amygdalothalamic and amygdalocortical projections in the monkey. The Journal of Comparative Neurology 502, 309–324.

Motzkin, J.C., Philippi, C.L., Wolf, R.C., Baskaya, M.K., and Koenigs, M. (2015). Ventromedial prefrontal cortex is critical for the regulation of amygdala activity in humans. Biological Psychiatry 77, 276–284.

Nummenmaa, L., and Saarimäki, H. (2017). Emotions as discrete patterns of systemic activity. Neuroscience Letters in press.

O&Reardon, J.P., and al, e. (2007). Efficacy and safety of transcranial magnetic stimulation in the acute treatment of major depression: a multisite randomized controlled trial. Biological Psychiatry 62, 1208–1216.

Okon-Singer, H., Hendler, T., Pessoa, L., and Shackman, A. (2015). The neurobiology of emotion-cognition interactions: fundamental questions and strategies for future research. Frontiers in Human Neuroscience 9.

Oya, H., Howard, M.A., Magnotta, V.A., Kruger, A., Griffiths, T.D., Lemieux, L., Carmichael, D.W., Petkov, C.I., Kawasaki, H., Kovach, C.K., et al. (2017). Mapping effective connectivity in the human brain with concurrent intracranial electrical stimulation and BOLD-fMRI. Journal of Neuroscience Methods 277, 101–112.

Oya, H., Kawasaki, H., Dahdaleh, N.S., Wemmie, J.A., and Howard Iii, M.A. (2009). Stereotactic Atlas-Based Depth Electrode Localization in the Human Amygdala. Stereotactic and Functional Neurosurgery 87, 219–228.

Pearl, J. (2009). Causality (New York: Cambridge University Press).

Pessoa, L. (2013). The Cognitive-Emotional Brain: From Interactions to Integration (Cambridge, MA: MIT Press).

Poldrack, R. (2015). Long-term neural and physiological phenotyping of a single human. Nature Communications 6, 8885.

Ramsey, J.D., Glymour, M., Sanchez-Romero, R., and Glymour, C. (2016). A million variables and more: the fast greedy equivalence search algorithm for learning high-dimensional graphical causal models, with an application to functional magnetic resonance images. International Journal of Data Science and Analytics doi:10.1007/s41060-016-0032-z.

Reichenbach, H. (1956). The Direction of Time (Berkeley, CA: University of California Press).

Robinson, E.C., Jbabdi, S., Glasser, M.F., Andersson, J.L., Burgess, G.C., Harms, M.P., Smith, S.M., Van Essen, D.C., and Jenkinson, M. (2014). MSM: a new flexible framework for multimodal surface matching. Neuroimage 100, 414–426.

Saarimäki, H., Gotsopoulos, A., Jääskeläinen, I.P., Lampinen, J., Vuilleumier, P., Hari, R., Sams, M., and Nummenmaa, L. (2015). Discrete neural signatures of basic emotions. Cerebral Cortex doi: 10.1093/cercor/bhv086.

Shackman, A.J., and Fox, A.S. (2016). Contributions of the central extended amygdala to fear and anxiety. The Journal of Neuroscience 36, 8050–8063.

Shackman, A.J., Fox, A.S., Lapate, R.C., and Davidson, R.J., eds. (2018). The nature of emotion: fundamental questions, 2nd edn (New York: Oxford University Press).

Smith, S.M. (2012). The future of fMRI connectivity. Neuroimage 62, 1257–1266.

Smith, S.M., Miller, K.L., Salimi-Khourshidi, G., Webster, M., Beckmann, C.F., Nichols, T.E., Ramsey, J.D., and Woolrich, M.W. (2011). Network modelling methods for fMRI. Neuroimage 54, 875–891.

Smith, S.M., Vidaurre, D., Beckmann, C.F., Glasser, M.F., Jenkinson, M., Miller, K.L., and Van Essen, D.C. (2013). Functional connectomics from resting-state fMRI. Trends in Cognitive Sciences 17, 666–682.

Spirtes, P., Glymour, C., and Scheines, R. (2000a). Prediction and Search, 2 edn (MIT Press).

Spirtes, P., Glymour, C.N., and Scheines, R. (2000b). Causation, prediction, and search (Cambridge, MA: MIT Press).

Sporns, O. (2013). The human connectome: origins and challenges. Neuroimage 80, 53–61.

Stokes, P.A., and Purdon, P.L. (2017). A study of problems encountered in Ganger causality analysis from a neuroscience perspective. PNAS 114, E7063–E7072.

Suthana, N., Haneef, Z., Stern, J., Mukamel, R., Behnke, E., Knowlton, B., and Fried, I. (2012). Memory enhancement and deep-brain stimulation of the entorhinal area. New England Journal of Medicine 366, 502–510.

Van Essen, D.C., Smith, S.M., Barch, D.M., Behrens, T.E., Yacoub, E., and Ugurbil, K. (2013). The WU-Minn Human Connectome Project: an overview. Neuroimage 80, 62–79.

Wager, T.D., Kang, J., Johnson, T.D., Nichols, T.E., Satpute, A.B., and Barrett, L.F. (2015). A Bayesian model of category-specific emotional brain responses. PloS Computational Biology DOI:10.1371/journal.pcbi.1004066.

Whalen, P., and Phelps, E.A. (2009). The human amygdala (New York: Oxford University Press).

Willie, J.T., Inman, C.S., Bass, D., Gross, R.E., and Hamann, S. (2016). Changes in autonomic arousal elicited by human amygdala stimulation are parameter-dependent. Paper presented at: Annual Meeting of the Society for Neuroscience.

Zhang, D., and Raichle, M.E. (2010). Disease and the brain&s dark energy. Nature Reviews Neurology 6, 15–28.

